# Comprehensive analysis of structural variants in breast cancer genomes using single molecule sequencing

**DOI:** 10.1101/847855

**Authors:** Sergey Aganezov, Sara Goodwin, Rachel Sherman, Fritz J. Sedlazeck, Gayatri Arun, Sonam Bhatia, Isac Lee, Melanie Kirsche, Robert Wappel, Melissa Kramer, Karen Kostroff, David L. Spector, Winston Timp, W. Richard McCombie, Michael C. Schatz

**Affiliations:** Johns Hopkins University, Baltimore, MD, 21211; Baylor College of Medicine, Houston, TX 77030; Cold Spring Harbor Laboratory, Cold Spring Harbor, NY 11724; Northwell Health, Lake Success, NY 11042

## Abstract

Improved identification of structural variants (SVs) in cancer can lead to more targeted and effective treatment options as well as advance our basic understanding of disease progression. We performed whole genome sequencing of the SKBR3 breast cancer cell-line and patient-derived tumor and normal organoids from two breast cancer patients using 10X/Illumina, PacBio, and Oxford Nanopore sequencing. We then inferred SVs and large-scale allele-specific copy number variants (CNVs) using an ensemble of methods. Our findings demonstrate that long-read sequencing allows for substantially more accurate and sensitive SV detection, with between 90% and 95% of variants supported by each long-read technology also supported by the other. We also report high accuracy for long-reads even at relatively low coverage (25x-30x). Furthermore, we inferred karyotypes from these data using our enhanced RCK algorithm to present a more accurate representation of the mutated cancer genomes, and find hundreds of variants affecting known cancer-related genes detectable only through long-read sequencing. These findings highlight the need for long-read sequencing of cancer genomes for the precise analysis of their genetic instability.

---

Somatic mutations that drive cancer development range across all genomic scales, from single nucleotide mutations through large-scale genome rearrangements, and have been observed in nearly all types of cancer at every stage of the disease progression^1^. Better detection, quantification, and reconciliation of mutation types in cancer samples can lead to a better understanding of disease progression and help improve existing and develop new, often patient-specific, therapeutic approaches for the disease^2^. Furthermore, improvements in detecting germline genetic variants in healthy cells can allow for better risk assessment of both hereditary and *de novo* mutations of various cancer types, leading to a more proactive rather than reactive cancer treatment approach^3^.

Our ability to detect genetic alterations has evolved over the last several decades of cancer genetics research. Prior to the completion of the human genome project, only a small handful of oncogenes or tumor suppressors were known^4,5^. Large-scale detection of cancer mutations began in around the year 2000 after the initial sequencing of the human genome using either microarrays^6,7^ or PCR amplification of known cancer-related genes from tumor and normal tissue and sequenced on low-throughput ABI capillary instruments^8^. In the late-2000s, the advent of Solexa, which later became Illumina, next-generation sequencing instruments dramatically accelerated the pace of discovery so that whole cancer genomes could be sequenced for the first time^9,10^. Since then, the dramatic improvements in the throughput and cost of whole-genome sequencing (WGS) and whole-exome sequencing (WES) over the past decade have made these technologies increasingly important in cancer studies, opening the door to widespread sequencing of patients, and the advancement of precision & personalized medicine. Within the Cancer Genome Atlas Project^11^ (TCGA), the International Cancer Genome Consortium^12^ (ICGC), and other large-scale efforts, several thousands of tumors have been sequenced using short-read Illumina sequencing across dozens of major cancer types. These studies have had a tremendous impact in cancer genomics, leading to the discovery, for example, of different signatures and mutation rates across cancer types, and new insights into the clonal structural and evolution of tumors^13–15^.

These results have substantially advanced our understanding of cancer susceptibility and progression, although the identification and understanding of the genetic alterations in cancer remains incomplete. A major contributor to our incomplete knowledge is that the known mutations have chiefly been detected using short-read Illumina sequencing^16^. This technology is very effective for identifying single nucleotide variants (SNVs) and large copy number variants (CNVs, especially those 100kb or larger), however, several studies have found it has poor accuracy for structural variant (SV) detection^17^. SVs are larger mutations, 50 bp or larger, where sequence is added, removed, or rearranged in the genome. Because of the short-read lengths, Illumina sequencing is difficult to map across SV breakpoints, especially insertions that are not present in the reference genome. Furthermore, SVs are frequently flanked by repetitive sequences so that the short-read sequence data frequently can not be unambiguously mapped back to its correct genomic position and *de novo* assembly techniques also fail to capture the novel sequences^18^. Consequently, short-read analysis approaches systematically fail to detect SVs, with false negative and false positive rates above 50%^19^. As a result, we are facing a major limitation with short-read sequencing studies of cancer where the field has systematically missed many important variants, potentially making it blind to entire classes of inherited genetic risk factors and blind to how SVs may mediate cancer progression and patient survival.

New long-read, single molecule sequencing technologies from Pacific Biosciences (PacBio) and Oxford Nanopore Technologies (ONT) have been shown to more reliably identify SVs with substantial improvements to both sensitivity and specificity. Reports by several groups have found a typical healthy human genome contains approximately twenty thousand SVs, and that they can be detected with 95% or greater sensitivity and specificity with long-reads^17,20,21^. These variants are especially important to accurately identify for somatic mutations that are not in linkage disequilibrium with any nearby SNVs. Long-reads can also improve the detection of SNVs and smaller insertion/deletion (indel) variants, especially in repetitive sequences and other sequences that are poorly resolved by short-reads^22,23^. Notably, 748 genes have been identified that are inaccessible to short-read sequencing^22^, including 193 medically-relevant genes with at least 1 exon that cannot be sequenced with short-reads, but are accessible to long-reads^24,25^. Long-reads also have improved power to resolve complex regions of the human genome, such as the highly variable major histocompatibility complex (MHC) or the lipoprotein(a) (LPA) gene sequence; and in some cases identified causative SVs underlying genetic disease that had been missed by short-reads^26,27^. Within cancer genetics, we previously published one of the first reports using PacBio long-read sequencing to study SVs in a cancer cell line genome and found that long-reads could detect tens of thousands of variants that had been missed by short-reads^17,28^. This included variants in known cancer genes such as HER2, APOBEC3B and CDH1, as well as dozens of novel gene fusions and other complex rearrangements that had substantially altered the expression and regulation of genes in the cell. Since this work, the cost and quality of 3^rd^-generation sequencing platforms make them more suitable than ever before in both academic and medical settings^21,29^, and thus require the improvement of existing and the development of new methods for mutation detection and analysis.

Addressing these questions, here we provide a comprehensive analysis of the SKBR3 breast cancer cell line and patient-derived organoids representing tumor and matching normal cells from two breast cancer patients sequenced with ONT, PacBio, and Illumina/10× 3^rd^-generation sequencing technologies. We identify and reconcile different types of large-scale genomic mutations in observed samples with an ensemble of methods, and highlight concordance and differences observed across different mutation inference methods and sequencing technologies. We demonstrate that long-read sequencing technologies outperform short-reads in SV detection with strong concordance between ONT and PacBio. Further, our integration of large-scale allele-specific CNVs and SVs into cancer genome karyotypes provides a more accurate representation of the observed mutated genomes. Notably, we observe hundreds of SVs, inferred exclusively with long-reads, that affect known cancer-related COSMIC^30^ census genes, and argue that long-read analysis of genetic variants plays a critically important in the area of cancer genomic instability. These results outline the advantages and limitations of various sequencing technologies when deployed for large-scale mutation detection in cancer genomes and provide a comprehensive biological and bioinformatics pipeline aimed at a more precise analysis of both patient-specific as well as overall genetic instability in cancer.

## Results

### Sample Identification and sequencing

Building on our previous work, we first evaluated the widely studied SKBR3 breast cancer cell line. SKBR3 is one of the most widely studied HER2+ breast cancer cell lines, with applications ranging from basic to preclinical research^31–34^. SKBR3 was chosen for this study due to its importance as a basic research model for cancer and because the SKBR3 genome contains many of the common features of cancer alterations including a number of gene fusions, oncogene amplifications, and extensive rearrangements. Notably, we previously sequenced this cell line using short-read paired-end Illumina and long-read PacBio sequencing^35^ allowing us to focus on 10X Genomics Linked Reads and Oxford Nanopore sequencing for this sample (**Figure 1** and Online Methods).

**Figure 1.**
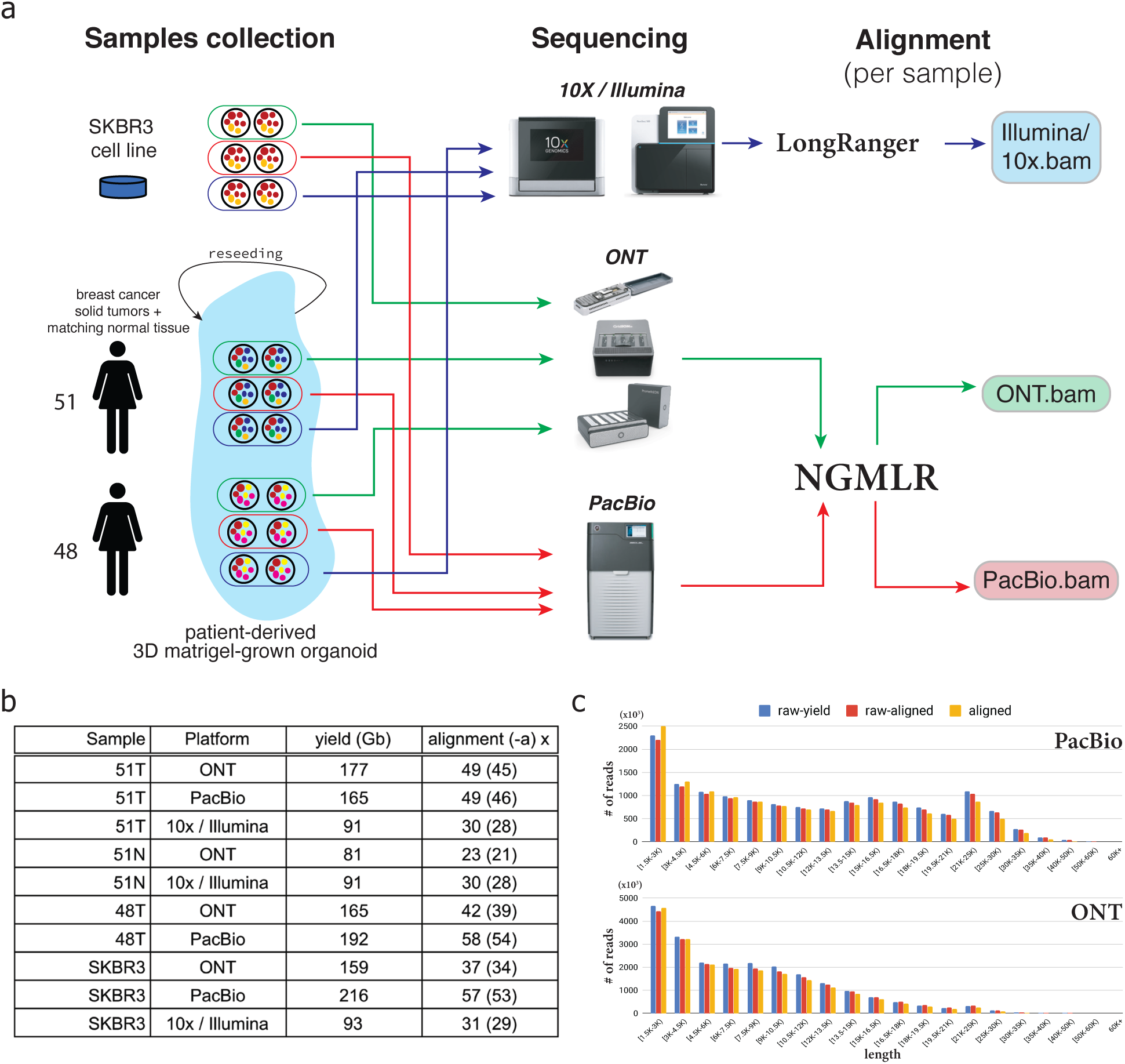
Sample collection, sequencing, and alignment pipeline and statistics overview. **a)** Biological data samples collection, sequencing, and alignment workflow for *SKBR3* breast cancer cell line and 3D Matrigel-grown organoids for solid breast cancer tumor tissues obtained from 2 female patients *51* and *48*. **b)** Yield and alignment coverage statistics for observed samples across WGS experiments various sequencing platforms. Suffixes *T* and *N* next to patients’ identifiers indicate tumor or matching normal tissue. Alignment values *x (y)* represent average read-depth *x* for aligned reads with *(y)* representing average read-depth when all unresolved *Ns* in the reference are also taken into consideration. **c)** Lengths distribution for reads of length *1*.*5+Kbp* from PacBio and ONT sequencing runs for patient *51. raw-yield* corresponds to lengths of raw sequenced reads, *raw-aligned* corresponds to lengths of raw read that had any alignment inferred for them and *aligned* corresponds to lengths of aligned parts of sequenced reads.

In addition, we sequenced tumor and normal patient-derived organoids from two breast cancer patients, here identified as patient 51 and patient 48, treated at Northwell Health and recruited in accordance with their Institutional Review Board Protocol (Online Methods). Patient-derived organoids are three-dimensional cultures of normal and cancer cells, propagated inside a basement-membrane extract matrix, overlaid with a growth-factor-rich growth medium tailored towards the origin of the individual tissue^36^. The organoids were generated from cells harvested from surgical specimens from the patients, and hence share the same genetic composition as the patient normal and tumor cells. Crucially, this allows for us to propagate the cells in a stable environment so that ample quantities of DNA were available for our three sequencing platforms (**Figure 1**). Furthermore, several studies have shown organoids are superior to standard 2D cell culture for recapitulating the molecular characteristics, physiology and treatment response of patient tissues^37^, allowing us to perform methylation analysis, RNAseq and other assays on the samples.

### Improved sensitivity and high concordance in structural variation inference with ONT and PacBio long-reads

Using these data, we then utilized an ensemble of methods to infer all types of SVs at least 50bp in size, including insertions, deletions, inversions, translocations, and duplications. For both ONT and PacBio datasets we used two state-of the art methods Sniffles^17^ and PBSV (https://github.com/PacificBiosciences/pbsv), and for Illumina/10X dataset we used 6 SV inference methods, with 3 (Lumpy^38^, Manta^39^, and SvABA^40^) designed for regular paired-end short Illumina reads, and 3 (NAIBR^41^, GrocSVS^42^, and LongRanger^43^) which also utilize the single-molecule 10X Genomics barcode information. We then iteratively merged SVs using the SURVIVOR^44^ package, first merging calls from all SV detection methods for each sequencing technology separately, and then merging across sequencing technologies to obtain sample-specific SV callsets (**Figure 2a**).

**Figure 2.**
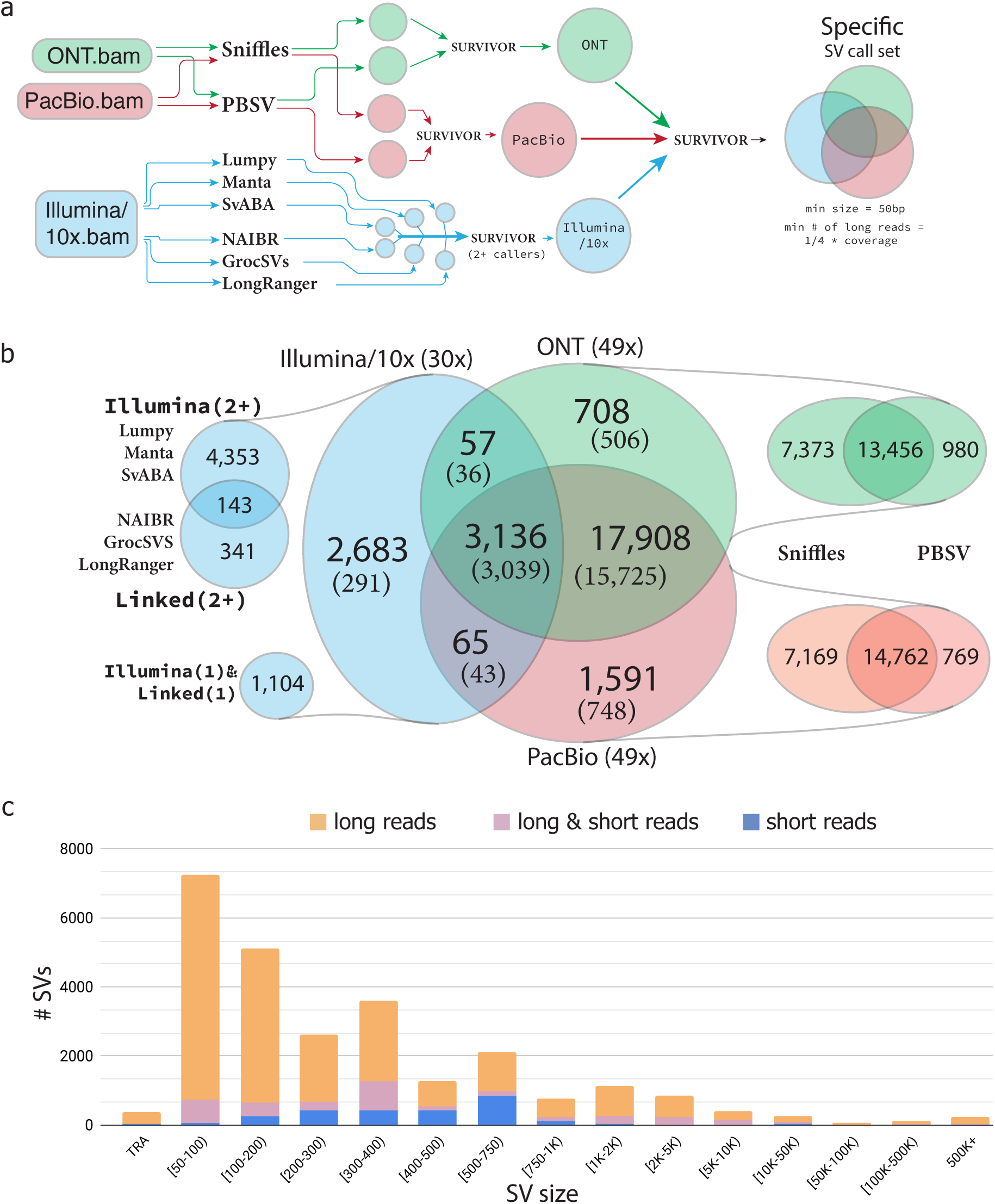
Structural variantions inference across Illumina/10X, ONT, and PacBio sequencing platforms for sample *51*. **a)** Ensemble worfklow for SV inference, with multiple methods and technologies used to infer SVs, subsequent merging of, first method-specific results, and then technology-specific results, with size and support restrictions applied. **b)** SV inference comparison across SVs inferred from *Platform (x)* sequencing exeriments, where *Platform* corresponds to sequencing technology, and *(x)* determines the average alignment read-depth coverage in the tumor sample. Methods-specific breakdown is provided for every sequencing technology. SVs detected in the normal sample are in parentheses. **c)** Size distribution for SVs in sample 51T with SVs being either exclusively inferred from either long-reads (either ONT, or PacBio, or both), or exclusively from Illumina/10x short-reads, or supported by both long and short-reads.

Since SVs inferred from paired-end short-reads are notorious for high rates of false positives^17,45,46^, for the Illumina/10X dataset we only considered SVs supported by at least 2 methods. To mitigate false positives in the long-read SV calls we only report SVs that were supported by at least one quarter of the average alignment read-depth in either ONT or PacBio datasets (also see **Coverage requirements** below). During the merging, we optimize parameters to minimize the effects of small thresholding issues, such as a variant present in 10 reads in one sample, and hence called as a variant, but only 9 reads in other, and hence not called (see Online Methods for details). Our results indicate a very strong concordance between SVs inferred with ONT and PacBio. Between 90% and 95% of variants called in at least one of the long-read data types were supported by both, with slightly lower concordance between PacBio-only calls (**Figure 2b** and **Supplementary Figure 2a,c**). We observe that while more than 50% of SVs inferred from short-read data were also identified by long-reads, the total quantity of SVs inferred from short-reads is approximately 4 times less than for either of the long-read-based inferences. We also demonstrate that across all SVs sizes, long-read based SV inference outperforms that of short-reads (**Figure 2c** and **Supplementary Figure 2b**).

For patient 51 for which we sequenced both the tumor and the matching normal cells we observed that 77% (20,388/26,148) of the SVs identified in the tumor sample were also identified in the matching normal sample (**Figure 2b**). A high fraction of SVs present both in the cancer and in the normal cells is expected since the cancer cells originate from normal tissue. Cancer cells, however, will generally acquire new mutations resulting in the addition of nearly 6,000 variants, although large deletions and loss-of-heterozygosity can potentially decrease the count of inherited SVs^47^. We also observe that for SVs called exclusively by short-reads only ∼11% (291/2,683) of SVs inferred in the tumor were also present in the matching normal cells. This is severalfold less than for SVs inferred both exclusively with long-reads (88%), and with both long and short-reads (97%), and we attribute this discrepancy to a high false positive rate in short-read SV inference.

To better understand the level of patient-specific and shared germline SVs, both in observed patients and the SKBR3 cancer cell-line, we compared SVs inferred with multiple sequencing technologies in the presented study to SVs identified in 15 healthy human genomes sequenced with PacBio long-reads presented in the recent study by Audano *et al*^20^. We find a high level of agreement between the SVs themselves and the distributions of their breakends locations identified in the cancer samples and the healthy samples (**Supplementary Figure 3**). We observe that 2,577 of the tumor-only SVs in patient 51 are present in other observed healthy samples and we thus hypothesize that many of them are actually present in the normal cells of patient 51, and the inability to infer them in normal cells stems from the lack of coverage in the ONT and the absence of PacBio long-read sequencing of the normal sample. This conjecture is supported by the comparison of SV types exclusively inferred with different long-read sequencing technologies, since the vast majority (1,806/2,577) are insertions, with ∼70% having lengths of 50-200 bp. More accurate basecalling and better SV-genotyping algorithms can help address such limitation in the future.

We further examined the distribution of SVs signature-based types across technology-specific subsets of inferred SVs in the analyzed cancer samples as well as in the healthy SV set (**Supplementary Figure 4**). We observe that in SVs identified exclusively with long-reads insertions and deletions comprise the largest fractions of ∼0.5 and ∼0.36 respectively. Duplications comprise only ∼0.06 of the long-read exclusive SVs callsets, while the inclusion of SVs inferred with even 2+ short-read SV callers increases that value severalfold to ∼0.13-0.16 in the multi-technology SVs callset, and we further observe that duplications constitute ∼0.71-0.93 of the 10X/Illumina-exclusive SVs. We further observe that both inversion and translocation SV signatures constitute similar fractions in both cancer and healthy SV sets in either short, long, or multi-technology SV sets.

Nanopore sequencing has the unique ability to identify cytosine methylation changes from DNA sequencing data without any additional library preparation^63^. Using this capability, we also examined methylation characteristics for the observed cancer and normal genomes (see Online Methods). As previously reported^48^, we observed genome-wide hypomethylation in the tumor samples relative to normal (**Supplementary Figure 10a,b**). While this hypomethylation occurs genome-wide in the cancer genomes, when genetic contexts are considered promoter regions stand out as an exception to this trend (**Supplementary Figure 10c,d)**. Furthermore, promoter regions that had SVs in them showed a slight enrichment for hypomethylation when compared to promoter regions without SVs (**Supplementary Figure 10e**). We also observed similar averaged methylation frequencies’ trends around transcription start sites (TSS) both in cancer and normal samples (**Supplementary Figure 10f**). We also identified several notable examples where SVs located within promoter regions of known COSMIC genes coincide with methylation changes between samples normal and tumor cells in patient 51: (i) insertion in PRDM1 gene coincides with hypomethylation of the respective promoter region in the tumor (**Supplementary Figure 10g**); (ii) duplication in the promoter region of the ZEB1 gene coincides with the increased methylation of the affected area in the tumor (**Supplementary Figure 10h**); (iii) insertion in the promoter region of USP6 gene coincides with the blocking of the TSS demethylation in tumor (**Supplementary Figure 10i**). These findings demonstrate that long-read ONT sequencing of DNA obtained from patient-derived organoids can allow for an integrative analysis of both SVs and methylation, both of which play an important role in cancer studies.

Additionally, RNA-seq gene expression data obtained from the tumor 51T and the matching normal 51N samples to investigate the impact of SVs on transcription. For this, we focused on differences in expression levels of those genes affected by SVs present in 51T but not present in 51N or fifteen other healthy samples sequenced with long-reads (see Online Methods for details). Overall, we see that duplications and deletions generally increase and decrease gene expression, respectively, especially when more than 50% of the gene sequence is impacted by an SV (**Supplementary Figure 9a**). While in some examples (**Supplementary Figure 9b**) we observe SVs’ link to gene expression change more clearly, there is a considerable range in the fold change of the expression levels so that we cannot conclude that the magnitude of expression changes is solely explained by the considered SVs. We note that SVs of different types that span genes do not always uniquely determine the copy number changes of the affected genomic regions due to the rearranged nature of underlying cancer chromosomes. For example, localized deletions within larger highly amplified regions can still show an overall increase in genomic copy number and increase in expression. These examples highlight an important, yet evidently non-exclusive, role that somatic SVs can play in tumor cells development and progression, and thus the importance of SV detection in cancer studies.

Overall, our results demonstrate that for comprehensive SV inference single-molecule long-read sequencing is essential, with ONT and PacBio producing highly similar results, thus providing validation of the long read variant calls. We also observe that a majority of SVs detected in tumor samples are also present in both matching normal cells as well as in the union set of SVs from healthy samples. These observations also underscore the need for *patient-specific reference genome* approach in the analysis of structural variants and their role in cancer origination and progression.

### Coverage requirements for accurate structural variation inference with long-reads

As the limitations for SV inference with long-read technologies are largely cost driven, we have measured how robust the SV inference with either ONT or PacBio reads is to lower sequencing coverage. For this, we downsampled our full coverage datasets to various coverage levels, with the lowest one set to 5x, and then compared SVs inferred on the downsampled datasets to the ground truth SV callsets from the original full coverage datasets. As with all variant callers, long-read variant callers report variants supported by a certain minimum number of reads although the higher error rate for long read potentially makes this analysis more challenging. We measured this effect by adjusting the minimum number of long-reads required to span (i.e., support) an SV for it to be classified as present from 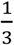 to 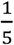 of the average read-depth coverage (**Figure 3a**). We found that for both ONT and PacBio reads the recall reaches a robust value of >0.8 and the precision reaches >0.9 with 24x to 30x coverage available (**Figure 3b** and **Supplementary Figure 5**). Both ONT and PacBio datasets also showed generally high consistency for minimum read supports, except for very low coverage datasets (<10x). These observations allow us to conclude that robust SV detection with single-molecule long-read sequencing is possible even at relatively low coverage levels of 25-30x average read-depth, with very similar results achievable with either ONT or PacBio long-reads.

**Figure 3.**
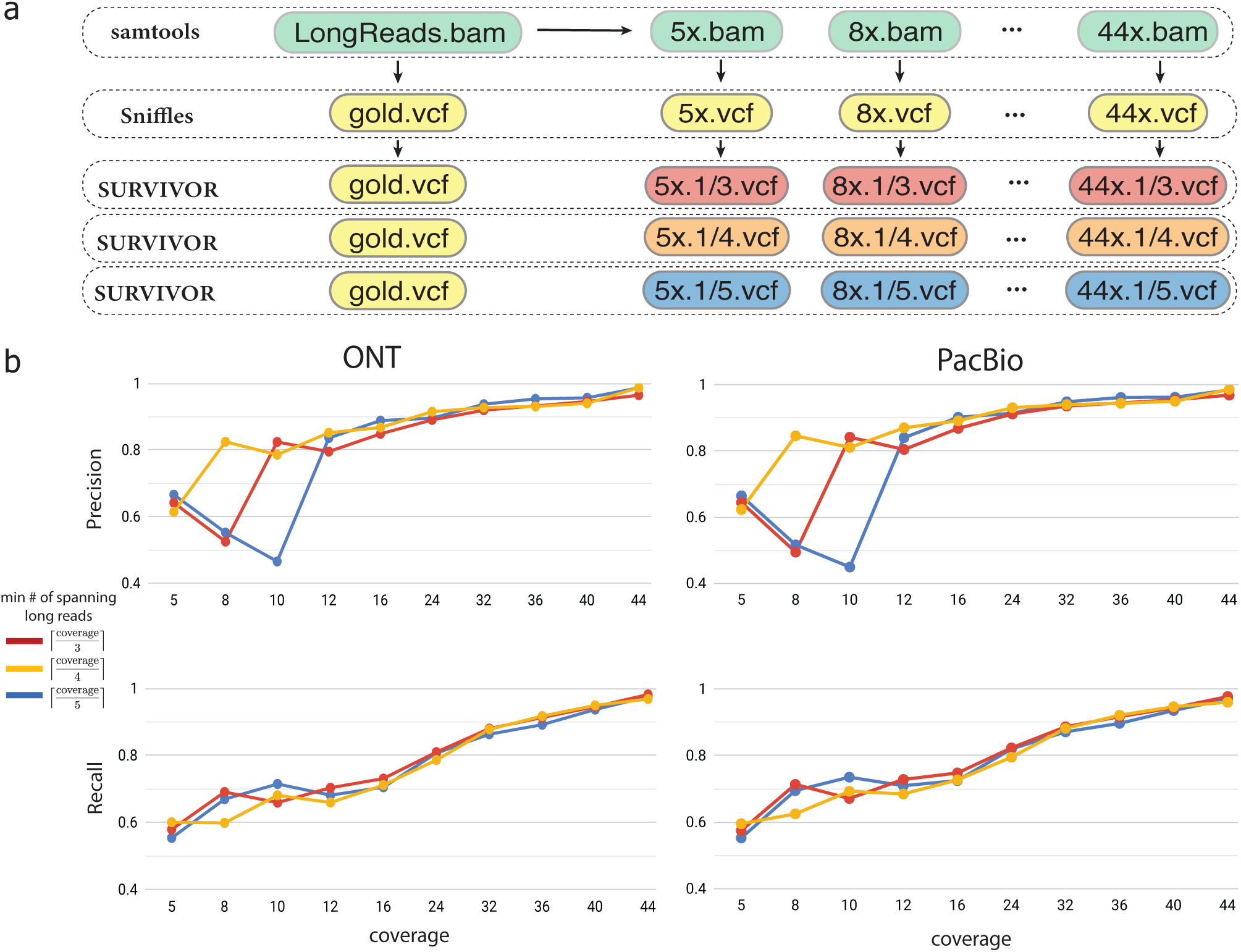
Structural variantions inference on downsampled long-read datasets. **a)** Workflow for downsampling full long-read dataset, and computing concordance between downsampled and full coverage datasets with distinct minimum fractional *x/y* read support for an SV to be considered. **b)** Precision and Recall for SVs inferred on downsampeld ONT and PacBio data for sample *51T*. SVs inferred on the full coverage dataset at the matching read support threshold are used as the ground truth.

### Cancer karyotype reconstruction

With the consensus SV callsets available, we then reconstructed the rearranged structure of the observed cancer genome for patient *51*, for whom both tumor and matching normal both short- and long-read sequencing are available. We utilized short-reads to infer large-scale allele-specific CNVs and inferred the homogeneous sample composition by using state-of-the-art methods TitanCNA^49^ and HATCHet^50^. These methods infer segment copy number profiles by observing changes in short-read coverage levels across large fragments and by using the germline Single Nucleotide Polymorphisms (SNPs) to infer copy numbers in the allele-specific fashion. While these methods provide a genome-wide view of large CNVs, the allele/haplotype information is lost across both adjacent and distant fragments, and smaller (<50Kbp) CNVs are also often missed.

To mitigate these limitations and to incorporate SV information into the analysis of rearranged cancer genomes we extended our recently proposed method RCK51 and utilized it to reconstruct haplotype-specific cancer karyotypes for patient 51 (see Online Methods). Briefly, RCK reconciles rearranged genomic segments, reference adjacencies, novel adjacencies (i.e., SVs) between segments’ extremities, and their copy numbers, and infers a haplotype-specific karyotype graph – or karyotype – which describes an alignment between the cancer and a healthy genome. RCK ensures that there exists a biologically plausible rearranged cancer genome with segment and adjacency copy number profiles determined by the inferred karyotype. RCK takes into account non-haploid nature of both normal and mutated genomes and also additionally supports long-read-informed haplotype constraints on the groups SVs breakends in the recovered karyotype (**Figure 4b**). In the new RCK v. 1.1 we added the support for long-read based haplotype constraints information for groups of SVs breakends, which helps to resolve ambiguities arising from equally plausible solutions in haplotype assignment for breakends during the karyotype inference process.

**Figure 4.**
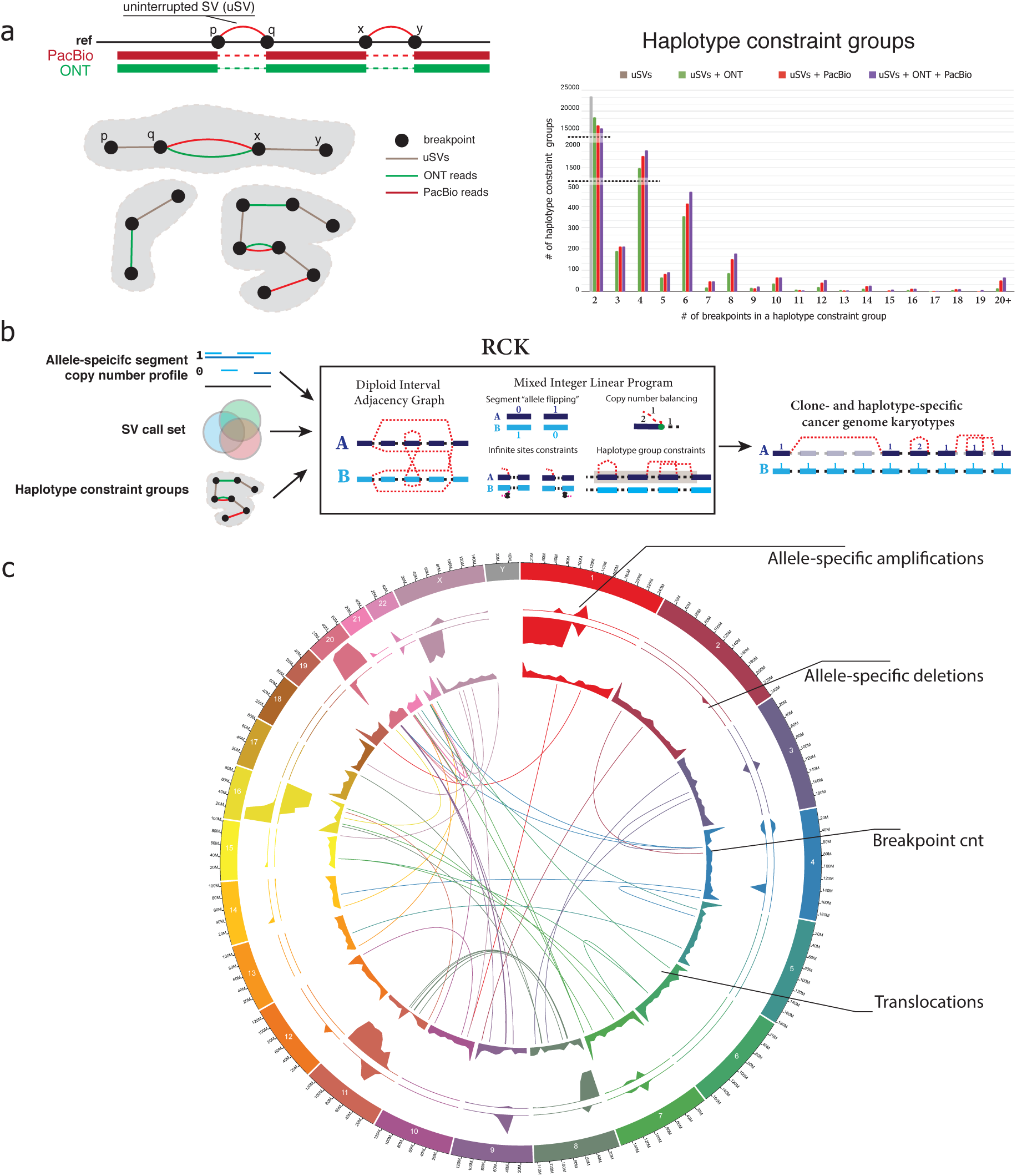
Cancer karyotype reconstruction. **a)**Haplotype constraint groups determined via uninterrupted SVs (uSVs) and long ONT and/or PacBio reads spanning multiple SVs. Statistics over the number of haplotype constraint groups inferred with only uSVs, and various combinations of uSVs and short/long-reads in patient 51. **b)** Workflow of the RCK method for reconstruction of haplotype-specific <u>c</u>ancer <u>k</u>aryotypes with allele-specific copy number profiles on large fragments, resolved SV callset, and inferred haplotype constraint groups. **c)** Circos plot of the karyotype inferred by RCK for patient 51 with HATCHet segment copy number (CN) input. Top two tracks corresponding to fractions *x/y* of the total length *x* of either amplified (CN ≥ 1) or deleted (CN = 0) fragments over the *y=5×10*^*6*^ long windows. Breakend track shows the total number (with 590 being the maximum value shown) of breakends inferred by RCK as being present. Translocation track shows inter-chromosomal SVs inferred by RCK as being present in the reconstructed katyorype.

We utilized both ONT and PacBio long-reads that spanned multiple SVs to introduce reference-haplotype-of-origin constraints, or haplotype constraint groups, (i.e., ensuring that grouped SVs breakends are assigned to the same haplotype) for the RCK karyotype inference (see Online Methods). Both ONT and PacBio demonstrated similar results in terms of grouping together multiple SVs breakends. We observed the expected prevalence of 2-breakend (i.e., single SV) haplotype constraint groups, but also identifying several hundreds of 6+-breakend groups (i.e., 3+ SVs), as well as few 20+ breakend groups where constraint information was obtained from multiple consecutively aligned long-reads and respective constraint groups joining via long-read overlaps (**Figure 4a**). The incorporation of haplotype constraint groups for SVs’ breakends reduces ambiguity in the cancer karyotype inference process and also helps to better assign haplotypes to allele-specific mutations, which have been shown to be important in previous cancer studies^52^.

We ran RCK with distinct input CNVs from both TitanCNA and HATCHet and with the same comprehensive SV callset, and obtained highly similar haplotype-specific karyotypes of the considered rearranged cancer genome (**Figure 4c** and **Supplementary Figure 6a**). We also observed that even though the CNV profiles of the inferred karyotypes were normalized by RCK to incorporate the input SVs, the resulting copy number profiles remained very similar to the input ones. This suggests that the initial large-scale CNV inference was compatible with the overall rearranged structure of the observed cancer genome and most of the alteration were necessary to refine the input CNV profiles in order to incorporate SV information and ensure the existence of the tenable rearranged chromosomes (**Supplementary Figure 5b**). We note that while up to a 10% of input SVs were allowed to be dismissed by RCK as either erroneous or not concurring with the input CNVs during the karyotype inference, we see very similar SVs breakend distribution across both the input SV callset, as well as in SVs utilized in the inferred karyotypes (**Supplementary Figure 7**). Overall, these results allowed us to study the impact of structural genetic variants on genes and other genomic features in much greater detail.

### Structural and copy number variants in COSMIC census genes

Using the reconstructed cancer karyotypes for 51T and inferred SVs for samples 48T and SKBR3 we considered how many of the reconciled structural and copy number variants affect known cancer-related genes cataloged in the COSMIC census gene set^53^. On average, more than twice as many SVs (622) affect COSMIC census genes as the genes being affected (237) in 51T (**Figure 5a**). This result held for both the initial SV callset and the refined SV set produced by RCK during the karyotype reconstructions. The majority (199/237) of the SV-affected COSMIC census genes in patient 51 were affected both the tumor and matching normal cells, and furthermore, a majority (466/622 in the initial SV callset and 428/568 in reconstructed karyotypes) of SVs affecting COSMIC census genes were also present in both the tumor and the matching normal cells. Long-read based SV inference identified five times as many COSMIC census genes affected by SVs and SVs affecting COSMIC census genes than was possible to infer with short-reads. Furthermore, the lack of concordance between SVs inferred exclusively with short-reads between the tumor and normal samples (6/79) provides additional evidence that the short-read SV calling is error-prone. In both patient 48 and the SKBR3 cell line we observed similar results (**Figure 5a**) with long-read SV inference outperforming short-read SV inference in both the number of COSMIC census genes affected, as well as the number of SVs affecting them.

**Figure 5.**
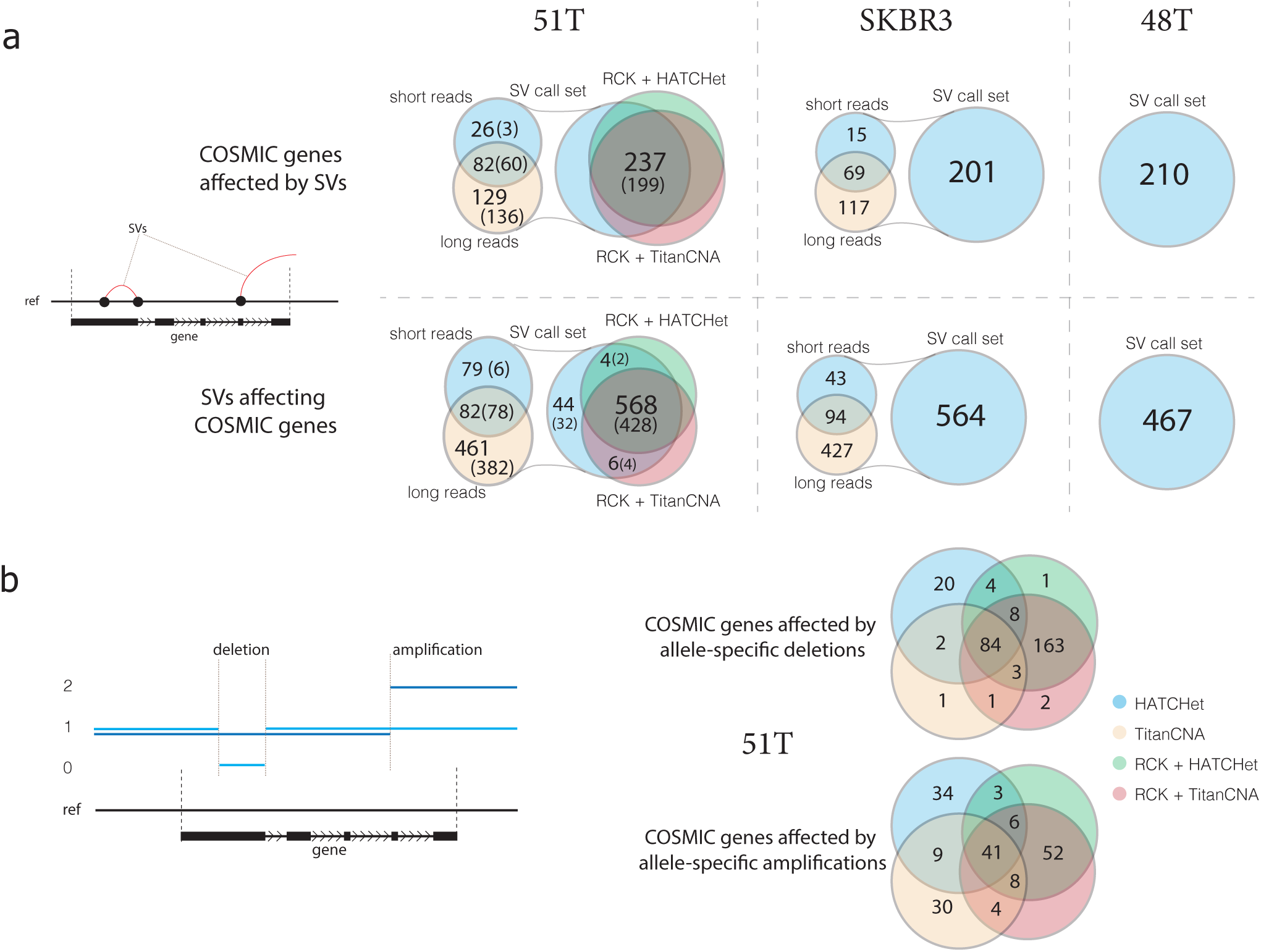
Structural and Copy Number Variants in COSMIC census genes. **a)** Comparison of the the number of COSMIC census genes being affected by SVs, as well as the number of SVs affecting COSMIC census genes, across inferred SV callset in 51T and N (parenthetical), SKBR3, and 48T, and SVs reported by RCK as being present in the karyptopes reconstructed with either HATCHet or TitanCNA copy number profiles in 51T. **b)** Comparison of the number of COSMIC census genes affected by either allele-specific deletions or amplifications between copy number profiles from HATCHet, RCK+HATCHet, TitanCNA, and RCK+TitanCNA in 51T.

To assess the population frequency of these variants, we genotyped identified SVs affecting COSMIC genes from the three analyzed cancer samples with Paragraph^54^ in the dataset of 2,504 short-read WGS samples from the recent re-sequencing of the 1000 genomes project (1KGP) samples^46^. Paragraph genotypes SVs by constructing localized sequence graphs containing the reference allele and the candidate SV allele and performs a localized realignment of paired-end short reads to the graph. The genotype is then determined based on the coverage of reads uniquely supporting the reference or variant allele breakpoints. Not all variants can be genotyped by Paragraph in all samples, resulting in no genotype call when support is ambiguous, so we consider only SVs that Paragraph was capable of genotyping in at least 1000 samples. We then summarize rare variants identified in <5%, <1%, and <0.1% of the overall observed samples (**Table 1**). We note that Paragraph v2.1 cannot genotype inversions, translocations, and large duplications, and thus we exclude such SVs from the genotyping analysis. SVs that were rarely present in 1KGP individuals (i.e., <0.01% frequency) were further filtered for variants which were not present in any of the 15 healthy genomes from the Audano *et al* study. We show that around 1/5 to 1/4 of the SVs we identified in COSMIC genes are genotyped at low frequency in the 1KGP individuals, and about half of these rarely genotyped SVs are also absent across all of the 15 healthy long-read genomes. These cancer variants found at low-frequency in a healthy population are thus the most likely candidates for cancer risk-factor-type mutations (Supplementary Table 1). These variants of interest are identified almost exclusively with long-reads, and although short-read genotyping can help determine population frequency, the ability of 15 long-read samples to additionally narrow the variants of interest further underscores the need for long-read sequenced genomes, both with healthy and disease phenotypes.

**Table 1.**
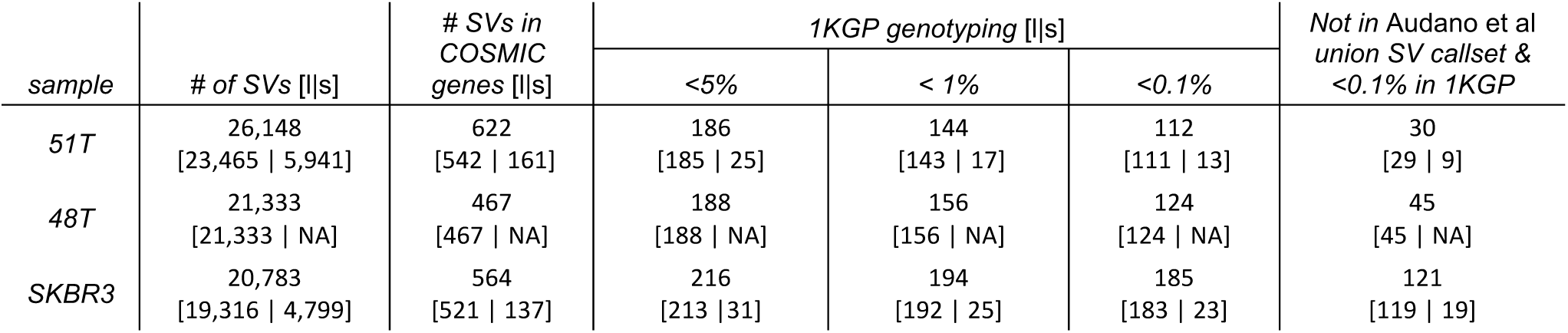
Genotyping of COSMIC genes’ affecting SVs in 1KGP and Audano *et al* datasets. For every observed tumor sample, we report the total number of identified SVs, the number of SVs directly affecting known COSMIC census genes, and the number of COSMIC genes’ affecting SVs that were successfully genotyped (i.e., called in at least 1000 samples) in 1KGP WGS short-read dataset with frequencies of <5%, <1%, and <0.1 % respectively. For the rarest (i.e., <0.1% in 1KGP) SVs report the number of such SVs that missing in the Audano *et al* union SV set. For every reported SVs count *x* we also show the numbers *[l|s]* of how many of SVs in *x* were inferred by long *(l)* or short *(s)* reads, respectively.

| sample | # of SVs [l\|s] | # SVs in COSMIC genes [l\|s] | 1KGP genotyping [l\|s] |  |  | Not in Audano et al union SV callset & <0.1% in 1KGP |
| --- | --- | --- | --- | --- | --- | --- |
|  |  |  | <5% | < 1% | <0.1% |  |
| 51T | 26,148<br>[23,465 \| 5,941] | 622<br>[542 \| 161] | 186<br>[185 \| 25] | 144<br>[143 \| 17] | 112<br>[111 \| 13] | 30<br>[29 \| 9] |
| 48T | 21,333<br>[21,333 \| NA] | 467<br>[467 \| NA] | 188<br>[188 \| NA] | 156<br>[156 \| NA] | 124<br>[124 \| NA] | 45<br>[45 \| NA] |
| SKBR3 | 20,783<br>[19,316 \| 4,799] | 564<br>[521 \| 137] | 216<br>[213 \| 31] | 194<br>[192 \| 25] | 185<br>[183 \| 23] | 121<br>[119 \| 19] |

We also observed in sample 51T a great concordance across COSMIC genes being affected by allele-specific CNVs as determined by copy number profiles in the inferred karyotypes as well as a strong overlap with the input large-scale CNVs (**Figure 5b**). The strong overlap between COSMIC genes affected by either of the two inferred karyotypes obtained from distinct CNV inputs, suggests that reconciliation of SVs an CNVs during the karyotype reconstruction process provides a way to mitigate possible noise and ambiguity that CNV-only inference methods may be faced with. This conjecture is also supported by a relatively low overlap over the subsets of the COSMIC genes that did not have CNVs affecting them in the RCK inferred karyotypes but were affected by CNVs in either TitanCNA or HATCHet CNV datasets.

Besides individual mutations affecting particular genes, the somatic evolution of cancer genomes is also known to be propagated by various complex rearrangements presence of which has been observed to have strong influence on the recovery prognosis of the patient^55,56^. In contrast with most often observed and studied rearrangements (e.g., deletions, insertions, tandem duplications, translocations) that require at most 2 double-stranded breakages, complex structural alterations may require 3+ breakages happening simultaneously. We identified, almost exclusively via SVs inferred with long-reads, several hundred complex rearrangements’ signatures in the initial SV calls set for samples 51T, SKBR3, and 48T, and in the reconstructed karyotypes for 51T, ranging from most frequently observed 3-breaks to a 7-break (**Supplementary Figure 8**). We note that not all complex k-breaks (k ≥ 3) produce reciprocal SVs, but without observing or reconstructing the actual somatic evolutionary history, reciprocal SVs breakends remain one of the best ways of identifying complex rearrangement events in cancer.

Four examples of genome rearrangements affecting COSMIC census genes in patient 51 are shown in **Figure 6**. We identified two insertions, which were missed by short-read SV inference, but were identified in both the ONT and PacBio datasets, in *BRCA1* and *CHEK2* breast cancer genes (**Figure 6a, b**). Both insertions have also been found exclusively with long-reads in the matching normal tissue, with the insertion in the *BRCA1* gene genotyped in <1% of 1KGP samples and present only in a single sample in the Audano *et al* dataset, suggesting a possible cancer-risk SV mutation, which would have been missed with widely-used next-generation whole genome sequencing analysis. In another example, we found multiple SVs present in *NOTCH1* and *ZNF331* COSMIC census genes (**Figure 6c, d**), which have been previously observed to play a significant role in tumor development^57,58^. Only one deletion in *NOTCH1* gene out of 6 observed SVs affecting *NOTCH1* and *ZNF331*, has been inferred from short-read data, while all 6 of the considered SVs were identified in both ONT and PacBio long-read datasets. The exon-spanning deletion in *ZNF331* present in both 51T and 51N samples was found in <1% of 1KGP samples but was identified in 3/15 samples in the Audano *et al* dataset. By utilizing long-reads that span multiple SVs at the same time we were able to assign all of the considered SVs in both *NOTCH1* and *ZNF331* genes to the respective reference haplotypes, with long-reads from both ONT and PacBio providing similar long-range haplotype constraint information. Assignment of multiple SVs to either the same or different haplotypes helps to better understand relationships between allele-specific genetic alterations, which has been observed^52^ to provide important information in determining the possible effects on the functional activity of the affected genes.

**Figure 6.**
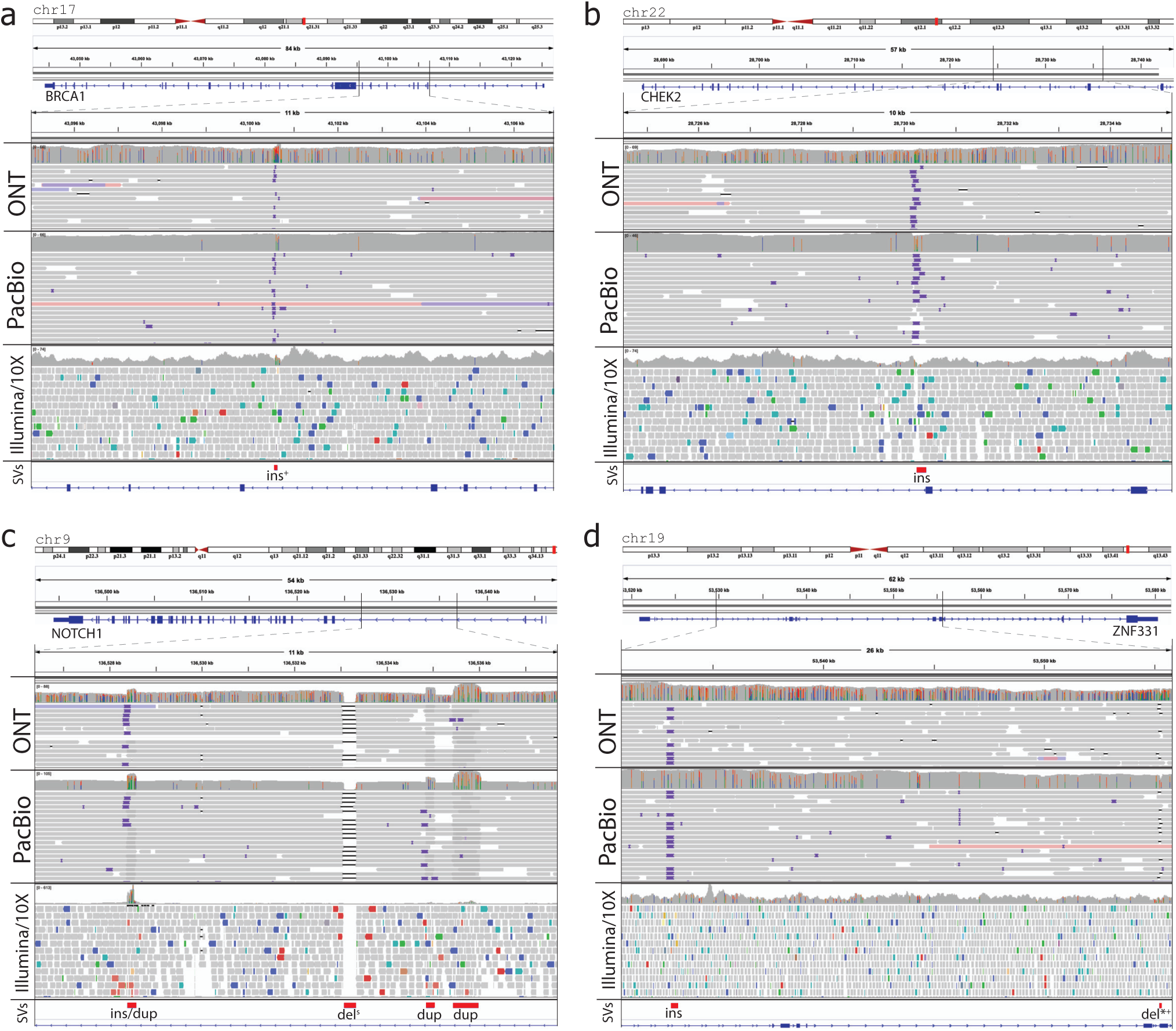
SVs identified in cancer-related COSMIC census genes in patient 51. All presented SVs are identified with both ONT and PacBio reads. Superscript marks *, +, and *s* indicate that marked SVs affect known exons, found as rare in 1KGP samples, and identified by short-read SV inference methods respectively. **a)** An insertion in the *BRCA1* gene identified in <1% of samples in 1KGP samples. **b)** An insertion in the *CHEK2* gene. **c)** An insertion/duplication, deletion, and two duplications in the *NOTCH1* gene, with deletion also found with short-reads. All 4 SVs belong to the same haplotype as indicated by multiple long (both ONT and PACBIO) reads spanning all of them at the same time. **d)** An insertion, and a deletion in the *ZNF331* gene, with the later deletion affecting an exon in the *NM_001317121* transcript, and gentyped in < 1% of 1KGP project samples. Both SVs belong to the same haplotype as indicated by long-reads spanning all of them at the same time.

Our utilization of inferred karyotypes with reconciled comprehensive SV and CNV callsets highlights the importance of incorporating various large-scale genomic variants in the comprehensive analysis of cancer genomes’ instability. We note that the abundance of long-read-exclusive SVs affecting COSMIC census genes both in the tumor and matching normal cells with some of them being genotyped only in a small number of samples from the 1KGP, and an even smaller subset being also absent across the union of long-read SVs in 15 healthy genomes from the Audano *et al* dataset, demonstrates the importance and utility of long-read sequencing as an avenue for cancer risk-factor analysis in healthy individuals as well as in the analysis of cancer drivers and its progression when studying rearranged tumor genomes.

## Discussion

In this study we presented a comprehensive analysis of three cancer genomes which we sequenced with Illumina/10X, ONT and PacBio sequencing technologies, and subsequently analyzed for large-scale (≥ 50bp) structural genomic mutations with an ensemble of methods. We observed how various SV and CNV inference methods compare to one another, and how SV inference results differ across considered sequencing technologies. We also demonstrated how SV and CNV mutations can be utilized together, reconciled, and integrated to infer a haplotype-specific cancer genome karyotype, which provides a refined view into the rearranged structure of observed cancer genomes.

Our findings demonstrate that current long-read sequencing technologies can be utilized in clinical settings to greatly improve genome-informed cancer risk assessment, analysis and treatment. We observe that while long-reads provide previously unprecedented resolution for SV detection, the sample preparation, sequencing, and analysis is on-par with that of similar short-read genome sequencing assays both in terms of complexity, time, and computational requirements. While costs are a major consideration for a technology to become widely applicable for patient care, we show that robust SV detection is possible at relatively low ∼30x average read-depth coverage with either ONT or PacBio long-read sequencing platforms. When applied at scale, costs for 30x coverage is below $1000 per sample for ONT PromethION and below $2,000 for PacBio Sequel II, which is highly comparable to ∼$800/$1,000 (Illumina/10X) for short-read sequencing.

In the presented study we observe the importance and power of integrating both SV and CNV signals into a comprehensive karyotype, which better describes the structural alterations in the observed mutated cancer genomes. As both SV and CNV callsets describe complementary measurements of the true underlying rearranged chromosomes, their integration allows for refinement of both large-scale CNVs as well as identification of spurious SV calls. We note that long-reads provide both unprecedented resolution in SVs inference as well as haplotype of origin constraints for groups of SVs and their breakends, which can be important when determining effects of multiple closely located SVs on the underlying functional sequences. Furthermore, while these cancer samples were homogeneous due to their cell-line/organoid-grown nature, direct patient cancer samples are often heterogeneous and comprised of multiple cancer clones with possibly distinct karyotypes^50,59^. For such samples, long-reads can provide valuable insight in assigning groups of SVs to particular clones and future long-read powered cancer studies can illuminate previously unseen aspects of clonal evolution in cancer.

We also note that as long-read sequencing technologies become more and more advanced it becomes possible to move away from a generic haploid human genome reference into an era of patient-specific reference sequences. We believe that future extension of the presented methodology can benefit from incorporating patient-specific diploid healthy genome structure as a starting point for mutation inference. We further underscore the importance of extending existing and developing new methods for multi-sample, time-series, and even multi-patient (in hereditary cases), integrative analysis of genetic instability that drives and propagates cancer development.

## Data and Code Availability

Sequencing data are in submission to dbGAP. The SV inference and comparison workflow is implemented with Snakemake^60^ v 5.5.4 and is available at github.com/aganezov/EnsembleSV. RCK v 1.1 utilized for cancer genome karyotype inference is available at github.com/aganezov/RCK.

## Acknowledgements

The authors would like to thank Benjamin J. Raphael, Rebecca Elyanow, and Simonne Zaccaria for thoughtful and productive discussions and their help in running the NAIBR and HATCHet methods. We would like to thank Gavin Ha for thoughtful and productive discussion and his help in running the TitanCNA method. This work was funded, in part, by the US National Science Foundation (NSF) grant DBI-1350041 (MCS); US National Institute of Health (NIH) grants R01-HG006677 (MCS) and 1R01HG009190 (WT); the Bill and Melinda Gates Foundation (MCS); and CSHL/Northwell Health (WRM,MCS,DLS), WT owns two patents currently licensed by Oxford Nanopore Technologies Limited. MCS and WT have received travel funding from Oxford Nanopore Technologies Limited.

