## Supplementary material for "Comprehensive analysis of structural variants in breast cancer genomes using single molecule sequencing": Online Methods

#### Patient-derived organoid culture.

Tumor resections from breast cancer patients along with adjacent normal tissue were collected from Northwell Health in accordance with Institutional Review Board protocol IRB-03-012 (TAP16-08). The collection of genomic and phenotypic data for this project was consistent with 45 CFR Part 46 (Protection of Human Subjects) and the NIH Genomic Data Sharing (GDS) Policy. Informed consent ensured that the de-identified materials collected, the models created, and data generated from them can be shared without exceptions with researchers in the scientific community. Patient-derived tumor and normal organoids were developed in accordance with a previously published protocol[36]. Briefly, the resected samples were manually cut into smaller pieces and treated with Collagenase IV at 37C. The samples were then manually broken down by pipetting into smaller fragments and seeded in a dome of matrigel. The organoids were grown in organoid culture media which contained 10% R-Spondin 1 conditioned media, 5nM Neuregulin 1 (Peprotech 100-03), 5ng/ml FGF7 (Peprotech 100-19), 20ng/ml FGF10 (Peprotech 100-26), 5ng/ml EGF (Peprotech AF-100-15), 100ng/ml Noggin (Peprotech 120-10C), 500nM A83-01 (Tocris 2939), 5uM Y-27632 (Abmole Y-27632), 1.2uM SB202190 (Sigma S7067), 1x B27 supplement (Gibco 17504-44), 1.25mM N-Acetylcysteine (Sigma A9165), 5mM Nicotinamide (Sigma N0636), 1x Glutamax (Invitrogen 12634-034), 10mM Hepes (Invitrogen 15630-056), 100U/ml Pen-Strep (Invitrogen 15140-122) 50ug/ml Primocin (Invitrogen ant-pm-1) in 1x Advanced DMEM-F12 (Invitrogen 12634-034) (Sachs et al., 2018). Organoids were passaged every 2-4 weeks using TrypLE™ (Thermo Fischer 12605028) to break down the organoids into smaller clusters of cells and re-plating them.

#### Organoid DNA and RNA extraction.

RNA was extracted using TRIzol® (Thermo Fischer 15596018) RNA extraction protocol. DNA was extracted by removing matrigel from organoids using ice cold PBS or TrypLE following by DNA extraction using Qiagen DNeasy Blood and Tissue kit (Qiagen 69504).

#### SKBR3 growth

SKBR3 cells were purchased from ATCC (ATCC HTB 30). Cells were grown in 5ml of McCoy 5A Medium (ATCC 30-2007) with 10% Fetal Bovine Serum ATCC 30-2020) and 1% penicillin/streptomycin (Sigma-Aldrich 11074440001). Cells were grown at 37℃ with 5% $CO_2$. To harvest cells, the media was removed and 2ml of Trypsin-EDTA 0.25% (Sigma 25200056) was added. Cells were allowed to sit at 37℃ for 10mins. The harvested cells were washed in PBS and either reseeded or used for DNA extraction.

#### Sample sequencing

DNA was sheared to >20kb via covaries G-tube (Covaris 520079). Oxford Nanopore DNA sequencing was carried out on a MinION or GridION device. Sheared DNA was prepared for sequencing using standard Oxford Nanopore methods. Briefly, Sheared DNA was repaired with the NEB FFPE repair module (NEB M6630L), ligated to Oxford Nanopore adapters (Oxford Nanopore SQK-LSK108) via the NEB blunt/TA master mix (NEB M0367L) and cleaned up with Ampure beads (Beckman Coulter A63881). The full volume of the prepared libraries was loaded on to a MinION R9.1 flow cell and run for 48 hours.

PacBio sequencing was carried out on a Pacific Biosciences Sequel I instrument using standard PacBio methods. Briefly, sheared DNA was prepared for sequencing via the SMRTbell template prep kit 1.0 (Pacific Biosciences 100-991-900). The prepared libraries were size selected on a Blue Pippin overnight with a 10-50kb range (Sage BUF7510). Libraries were loaded for sequencing on a 1M SMRTcell (Pacific Biosciences 101-531-000) with a concentration of 4-10pM with diffusion loading and 10 hour movies.

**Read alignment**
All read alignments were performed against latest human genome reference GRCh38[61]. ONT and PacBio long-reads were aligned with NGMLR[17] v 0.2.7. 10X/Illumina short-reads were aligned with LongRanger[43] v 2.1.6 pipeline. Only major chromosomes 1-22, X were considered for the alignment and subsequent structural analysis. Alignment coverage was computed with samtools[62] v 1.9 depth command both with and without -a flag and computes an average of the per-base coverage values. For long-reads *raw-yield* lengths' distribution considers sequenced lengths of all reads. *Raw-aligned* lengths' distribution considers sequenced lengths reads that have at least some part(s) of them aligned to the reference. *Aligned* lengths' distribution considers lengths of aligned part(s) of sequenced reads.

**SV inference workflow.**
For both ONT and PacBio long-reads we used Sniffles v 1.0.11 and PBSV v 2.2.0 for SV inference. For Sniffles the minimum number of reads supporting SV was set to 2, and the minimum SV size was set to 30bp, although the final variant calls were a more stringent subset of these requiring higher read support and larger sizes. PBSV was run with default settings. For Illumina/10X reads we utilized, SVaBA v FH134, Lumpy v 0.2.13, Manta v 1.5.0, GROC-SVs v 0.2.5, NAIBR (version determined by 15eba96 commit GitHub master branch), and LongRanger v 2.1.6. All short-read SV callers were run with recommended settings. Some SV inference methods produced more than a single SV callset (usually with SVs segregated by size), which we subsequently concatenated into method-specific SV callsets. For example, for SVaBA we concatenated *indel* and *sv* SV callsets LongRanger we concatenated the *dels* and *large_svs* SV callsets,

For every sequencing technology the SVs produced by all callers were merged together with the SURVIVOR[44] v 1.0.6 software package into a *ONT*, *PacBio*, and *Ilumina/10X* technology-specific SV callsets. SURVIVOR merge was run with maximum distance between SVs set to 1000 and minimum SV size set to 30. SV types were not taken into account during the SURVIVOR merging as different methods may assign different types to the same inferred SV based on the respective method's terminology, but strand/orientation was required to match.

For SVs inferred on short-reads we removed any method-exclusive SVs (i.e., supported by only 1 out of 6 methods) and retained any SVs that had at least 2 methods inferring them. This was done in order to mitigate the well-known false positive problem that exists if the area of SV inference with short-read data.

To ensure consistency when comparing against 15 healthy genomes, sequenced with PacBio and reported in Audano *et al*[20], we performed alignment and variant calling on the 15 samples with the same pipeline described above. Raw reads from all 15 genomes were downloaded and aligned with NGMLR to the main chromosomes of GRCh38 with the -x pacbio setting. Structural variants were then called on each sample with Sniffles. As all samples were reported in Audano *et al* as above 40x coverage, we set Sniffles to require a minimum read support of 10 reads. A minimum SV size of 30bp was used. Comparisons against the SVs in these 15 genomes were performed with SURVIVOR merge with a maximum distance of 1000, type and strand considered, and a minimum size of 30, with thresholding performed post merging to examine only variants of at least 50bp, as described below.

**Comparison of SVs inferred with different sequencing technologies.**
ONT, PacBio, and Illumina/10X technology-specific SV callsets were subsequently merged together with the SURVIVOR package into a sample-specific *sensitive SV callset.* SV types were not taken into account for the same purpose as was described for the technology-specific merging procedure, minimum size for SVs to be considered was set to 30, maximum distance between SVs was set to 1000.

We further removed from the sensitive SV callset any of the SVs shorter than 50bp in order to focus only on large-scale rearrangements. This filtration was done after the merging of technology-specific SV callsets, rather than before, in order to mitigate thresholding issues that may have arisen if cases when the same underlying SV, which, due to possible noise in data, had a length of 49bp in one callset, and 51bp in another, would have the 49bp instance removed before it could have been merged with the 51bp instance, producing a 50bp-long merged SV, which would be retained.

To mitigate the relatively high per-basepair error rate in long-reads and its possible effect on false positive calls in long-read-exclusive (either from ONT, or PacBio, or both) SVs we removed long-read exclusive SVs for which then number of long-reads supporting them was less than a quarter of average read-depth in both ONT and PacBio datasets. After length and long-read-support filtration the we obtained the *specific SV callset* on which the agreement and discordance in SVs inference between sequencing technologies and methods was analyzed.

**Methylation analysis**
For calling methylation on ONT sequencing data, we used nanopolish[63] call-methylation module. A threshold of 2.5 was used to filter out ambiguous methylation calls. After aggregating methylation calls at each site, we smoothed the raw methylation frequencies using BSmooth function from Bioconductor R package bsseq[64].

Briefly, we first choose a window such that at least 50 CpG sites and 500 bps of region are covered for each locus $l_j$. Assuming that 1) the methylation frequency $f(l_j)$ follows a binomial distribution and 2) $\log\left(\frac{f(l_j)}{1-f(l_j)}\right)$ is approximated by a second degree polynomial, we fit a weighted generalized linear model inside each window. The weights are inversely proportional to the standard errors of the per-site measurements, and a tricube kernel is used in relation to the distance from the locus $l_j$ . For all subsequent analysis, we applied a coverage filter, removing data points on loci where the total number of calls were less than 5 in any of the samples.

For global comparisons of genomic context methylation, we used genomic contexts as determined by Ensembl[65] gene annotations and regulatory feature sets. For each region in the set, average methylation was calculated by dividing the sum of methylated calls by total calls.

**RNA-seq expression analysis**
RNA-seq libraries were prepared using Illumina TruSeq RNA Library prep kit v2 (RS-122-2001) and sequenced as 75bp paired-end. The reads were aligned using STAR-aligner[66] v 2.5.3a. The built-in gene counts option was used to count raw reads using gencode v27 gtf reference file. The counts files were exported into R v3.5.1 and normalized using DeSeq2[67] v1.22.2. Normalized counts were used to calculate log2 fold change of tumor versus normal samples.

SVs that were supported by long-read sequencing were used for this analysis. Structural variants overlapping genes was determined using the bedtools[68] `intersect –wo` command with gencode v27 gtf file as a reference. The graphs were plotted using ggplot2[69] v3.2.1. Percent overlap was calculated by dividing the number of overlapping base pairs with the total length of the gene.

**Downsampling and SV inference.**
We designed and implemented the downsampling workflow to analyze the robustness of long-read-informed SV inference at various read depth coverages. Every full coverage long-read ONT or PacBio alignment dataset *reads.bam* was downsampled with samtools v1.9 command *view -s x.y reads.bam*, where *x* determines the seed for randomize alignments selection to be included in the produced downsampled alignment, and $y$ determines the fraction of the read alignments from the initial dataset *reads.bam* to be selected.

For sample 51T the downsample coverage levels were set to $C = [5, 8, 10, 12, 16, 20, 24, 32, 38, 44]$x for both ONT and PacBio, for sample 48T coverage levels were set at $C = [5, 8, 10, 12, 16, 24, 32, 36, 40]$x for both ONT and PacBio, and for sample SKBR3 downsample coverage levels were set at $C = [5, 10, 16, 20, 24, 28, 32]$x or ONT, and additional coverage levels at $C = [40, 48, 52]$x were set for PacBio, as the PacBio dataset for SKBR3 had higher full coverage than the ONT one.

For both ONT and PacBio for every downsampled target coverage level we generated 3 distinct downsample read alignments datasets with different random seed values. SV inference on downsampled alignment datasets was carried with Sniffles v 1.0.11 with the minimum number of reads required to support an SV was set to 2, and a minimum SV size was set to 30. As previously described, to mitigate a relatively high per-basepair error rate in long-reads several reads are required to span an SV for it to be considered true. We observed how SV inference was affected by this parameter by considering various fraction $f \in [\frac{1}{3}, \frac{1}{4}, \frac{1}{5}]$ of an average downsample-dataset-specific read depth coverage as a threshold for the minimum number of reads required to span an SV. We then generated distinct $f$-SVs callsets by removing all SVs that were supported by less than a fraction $f$ of reads.

We then compared the technology-specific $f$-SVs callsets for every downsample coverage level $c \in C$ with the gold standard (i.e., SVs callset on full coverage dataset, using the matching read support threshold) with SURVIVOR and averaged the precision and recall results over 3 randomly created downsampled datasets for every coverage target level $c$.

**Somatic evolutionary model.**
We employ RCK‘s large-scale rearrangement somatic evolutionary model that describes extents and limitations for the organizations of the rearranged cancer genome(s). We briefly outline its main concepts below. A detailed description can be found in the original manuscript[51].

We assume that all the mutated cancer genomes evolve from a diploid reference genome $R$. Every chromosome in $R$ is present in two homologous copies, which we label $A$ and $B$ respectively. A segment $s_H = C: [s^t, s^h]$, where $H \in \{A, B\}$ is a contiguous part of the chromosome $C_H$, and its endpoints $s^t$, $s^h$ that determine the *tail* and the *head* of the segment are called *extremities*. We may omit chromosomal name in segment's signature, when obvious and/or irrelevant, depending on the context. Segments are labeled $1, \ldots, m$ throught the reference genome's chromosomes. Segments $(s_H, (s+1)_H)$ that are sequential on some chromosome determine an *adjacency* $\{s_H^h, (s+1)_H^t\}$. We denote by $\mathcal{A}(G)$ a set of adjacencies present in genome $G$, and we denote by $\mathcal{A}_N(G)$ a set of *novel adjacencies* (i.e., not present in the reference) present in genome $G$. We will omit the *tail* and *head* superscripts on extremities involved in adjacencies, when orientation is not important, just retaining the coordinates.

Outermost segments' extremities on every chromosome are called *telomeres*, and we denote by $\mathcal{T}(G)$ a set of telomeres in genome $G$.

We depict every large-scale rearrangement as a collection of double-strand breakages that destroy adjacencies, with possible amplification and/or loss of involved segments, and subsequent ligation of involved segments' extremities, which introduces novel adjacencies. We assume that somatic large-scale evolutionary history of observed cancer genomes complies with the Infinite Sites (IS) assumption, under which the same genomic coordinate (i.e., the same adjacency) on either $A$ or $B$ haplotype can be directly involved in at most one, however complex, genome rearrangement's breakage with subsequent ligation. We also assume that in rearranged genomes chromosomal telomeres are inherited from the reference, or more formally, $\mathcal{T}(G) \subseteq \mathcal{T}(R)$, as telomere genomic sequences play an important role in cells life cycles and they are required to contain specific sequences for the respective molecule replication to complete correctly.

**Haplotype constraint groups.**

For a genome $G$ the inferred SVs correspond to set $\widetilde{\mathcal{A}_N}(G)$ of unlabeled novel adjacencies (i.e., for every novel adjacency $a = \{p_F^x, q_D^y\} \in \mathcal{A}_N(G)$, where $F, D \in \{A, B\}$, and $x, y \in \{t, h\}$, we measure its unlabeled version $\tilde{a} = \{p^x, q^y\}$, or an SV, which is missing haplotype labels). Under the IS assumption every measured unlabeled novel adjacency $\tilde{a} = \{p^x, q^y\}$ has a unique haplotype-specific counterpart $a \in \mathcal{A}_N(G)$ (i.e., unique haplotype labels $F, D \in \{A, B\}$ such that $a = \{p_F^x, q_H^y\} \in \mathcal{A}_N(G)$). We also note, that we allow cases when multiple distinct measured SVs ($a = \{p, x\}, b = \{p, y\}, \dots$) involve the same unlabeled extremity $p$ and assume that all of underlying true SVs involve the same labeled segment's extremity on one of the two haplotypes.

We call a set $P = \{p, q, \dots\}$ of unlabeled extremities, or breakends, involved in measured SVs a *haplotype constraint group* if for every haplotype-specific version of the SVs involving extremities from $P$, the involved haplotype-specific extremities (e.g., $p_H, q_H$) belong to the same haplotype $H \in \{A, B\}$. We now describe how we obtain a set $\mathcal{P}$ of haplotype-constraint groups from the measurement data, as we extend the previous version of the RCK framework in which haplotype constraint groups were only determined for pairs of reciprocal (i.e., adjacent in the reference) breakends.

Given a set $\widetilde{\mathcal{A}_N}(G)$, of unlabeled novel adjacencies, or SVs, we call an inter-chromosomal SV $a = \{p, q\} \in \widetilde{\mathcal{A}_N}(G)$ *uninterrupted* (*uSV*) if $|q - p| < 5000$ and no other SV has a breakend overlapping with $a$, or, more formally, there does not exist an SV $b = \{u, v\} \in \widetilde{\mathcal{A}_N}(G)$, such that either $p \leq u \leq q$, or $p \leq v \leq q$, or both. We assume that every uninterrupted SV $a = \{p, q\}$ determines a haplotype-constraint group $P_a = \{p, q\}$, as the opposite will correspond to an unlikely event of 2+ double-strand breakages involving homologous copies of the same chromosome with breakage coordinates located very close to one another.

For a long-read $r$ that spans a set $S_r$ of SVs we consider the ordered sequence $O(S_r)$ of the SVs from $S_r$ and an ordered sequence $O(B(S_r))$ of breakends $B(S_r)$ as determined by $r$'s traversal of SVs in $S_r$. We note that since every SV determines a pair $\{p, q\}$ of breakends we naturally have $|O(B(S_r))| \equiv 0 (mod\ 2)$. For every consecutive pair $a, b$ of SVs from $O(S_r)$ let us observe the last breakend $p$ of $a$ and the first breakend $q$ of $b$ as determined by $O(B(S_r))$. Alternatively, we can say, that we observe every $2i^{\text{th}}$ and $(2i + 1)^{\text{st}}$ elements $p$ and $q$ of $O(B(S_r))$, where $i = \overline{1, |O(S_r)|}$. Since both SVs $a$ and $b$ are spanned by the same read, that means that there exists a segment $s = \{p, q\}$ in the sequenced cancer genome, which was not altered by any rearrangements (as otherwise there would be another SV and another breakends between p and q). Under the IS assumption the long-read $r$ a pair $\{p, q\}$ of breakends determines a haplotype constraint group $\{p, q\}$.

To obtain the set $\mathcal{P}$ of haplotype constraint groups for a set $\widetilde{\mathcal{A}_N}(G)$ of measured SVs we build a haplotype constraint graph $G_{\mathcal{P}} = (V, E)$. The set $V = \{p, q | a = \{p, q\} \in \widetilde{\mathcal{A}_N}(G)\}$ of vertices is determined by breakends involved in SVs from $\widetilde{\mathcal{A}_N}(G)$. The set $E$ of edges is constructed by adding edges $\{p, q\}$ that are determined by two-breakend haplotype constraint groups, which are either coming from reciprocal SVs' breakends, determined by uSVs, or inferred from long-reads spanning multiple SVs. We note that if there exist two haplotype constraint groups $P_1$ and $P_2$ such that $P_1 \cap P_2 \neq \emptyset$, then the set $P = P_1 \cup P_2$ is also a haplotype constraint group. The desired set $\mathcal{P}$ then corresponds to connected components of size (i.e., number of vertices) greater than 1 in $G_{\mathcal{P}}$.

**Large-scale CNV inference.**
To measure large-scale copy number variations (CNVs) we utilized Illumina/10X short-read sequencing datasets from both the tumor and the matching normal cells. We used TitanCNA and HATCHet CNV-inference methods, which produce clone- and allele-specific segment copy number profiles, or more formally, for a tumor sample $s = (G_1, G_2, \dots, G_n)$ comprised of $n$ genomes they produce what can be described as two matrices $(\widehat{\boldsymbol{C}}, \breve{\boldsymbol{C}}) = \left(\left[\left[\widehat{c_{i,j}}\right]\right], \left[\left[\widetilde{c_{i,j}}\right]\right]\right)$, where $\left(\widehat{c_{i,j}}, \widetilde{c_{i,j}}\right) \in \mathbb{N}^2_{\geq 0}$ determine allele-specific integer copy numbers for segment $j$ in genome $G_i$. In the considered patient 51 case both HATCHet and TitanCNA methods have inferred a homogeneous tumor composition with respect to cancer clones (i.e., a unique allele-specific CNV profile coming from a mutated cancer genome, with some admixture of normal cells, can explain the observed read-depth alterations for reads coming from the tumor sample 51T). We then treat the output of both HATCHet and TitanCNA as a pair of vectors ($\hat{\boldsymbol{c}} = [\widehat{c_1}, \widehat{c_2}, \dots, \widehat{c_m}], \breve{\boldsymbol{c}} = [\breve{c_1}, \breve{c_2}, \dots, \widetilde{c_m}]$), where $\left(\widehat{c_j}, \breve{c_j}\right) \in \mathbb{N}^2_{\geq 0}$ determine allele-specific copy numbers for segment $j$ in the cancer sole clone in the observed tumor sample 51T. Both methods were run with recommended settings.

We note that for every segment $j$ we are missing the true haplotype labels on the measured copy numbers, so rather than having true haplotype-specific segment copy numbers $(a_j, b_j) \in \mathbb{N}^2_{\geq 0}$, where $a_j$ and $b_j$ determines copy numbers of segments $j_A$ and $j_B$ in the mutated cancer genome, and thus, at best (assuming no errors in measurements) we have $\left\{\widehat{c_j}, \breve{c_j}\right\} = \left\{a_j, b_j\right\}$ (i.e., correct copy numbers, but missing order, or haplotype assignments). When considering possible errors in the measured copy number values we take into account that both methods infer CNV profiles on rather large ($\geq 50$Kbp) segments, which will miss any smaller copy number variations (e.g., small deletions or duplications), and the specific boundaries of large CNVs.

**Reconstruction of haplotype-specific karyotypes.**
In order to combine the SV and CNV mutation inference we utilized our RCK method which integrates both SV and CNV mutations together and infers the underlying clone- and haplotype-specific *karyotype graph* or simply *karyotype*. A detailed description of RCK can be found in the manuscript[51], and here we just provide a brief outline.

With the input set $\widetilde{\mathcal{A}_N}$ of SVs and allele-specific CNV profile $\widetilde{\boldsymbol{C}} = \{ (\hat{\boldsymbol{c}} = [\widehat{c_1}, \widehat{c_2}, \dots, \widehat{c_m}], \breve{\boldsymbol{c}} = [\breve{c_1}, \breve{c_2}, \dots, \widetilde{c_m}])$ RCK first constructs the set $S$ of segments such that every SV's breakend, or boundary of the fragment in CNV profile corresponds to the extremity of a segment from $S$. We refined the input CNV profile to represent copy number of refined segments, rather than input fragments, with copy number values for every segment $j$ being inherited from the respective spanning input fragment.

Then, RCK constructs the Diploid Interval Adjacency Graph (DIAG) $G = (V, E)$, where the set $V = \{j^t, j^h \mid j \in S\}$ of vertices is determined by the extremities from segments in set $S$, and the set $E = E_s \cup E_{\mathcal{A}}$ of edges is comprised two types of edges: (i) segment edges $E_s = \left\{\{j_H^t, j_H^h\} \mid j \in S; H \in \{A, B\}\right\}$, determined by haplotype specific segments from $S$, and (ii) adjacency edges $E_{\mathcal{A}} = \{a_R \mid a_r \in \mathcal{A}(R)\} \cup \left\{\{p_F^x, q_D^y\} \,\middle|\, \{p^x, q^y\} \in \widetilde{\mathcal{A}_N}; F, D \in \{A, B\}\right\}$. Adjacency edges determine all possible haplotype-specific transitions between segments' extremities in the observed cancer genome, which can be either inherited form the diploid reference $R$ or measured (with haplotype label loss) into $\widetilde{\mathcal{A}_N}$.

A karyotype graph is determined as DIAG $G$ with edge multiplicity function $\mu: E \to \mathbb{N}_{\geq 0}$ , where multiplicity $\mu(e)$ of the edge $e \in E$ determines how many times a respective segment/adjacency is present in the observed caner genome. For an underlying rearranged genome $G$ to exist and be described by a karyotype graph $G_\mu = (V, E, \mu)$ is it necessary and sufficient for the following two equations to be true:

$$\forall\, v \in V \setminus \mathcal{T}(G): \;\; \mu\big(e_s(v)\big) = \textstyle\sum_{e \in E_{\mathcal{A}}(v)} l(e) \cdot \mu(e),$$
$$\forall\, v \in \mathcal{T}(G): \mu\big(e_s(v)\big) \geq \textstyle\sum_{e \in E_{\mathcal{A}}(v)} l(e) \cdot \mu(e),$$

where $e_S(v) \in E_s$ is a segment edge incident to $v$, $E_{\mathcal{A}}(v) \subseteq E_{\mathcal{A}}$ are all of the adjacency edges incident to $v$, and $l(e): E \to \{1,2\}$ is an auxiliary function that output 2 if the edge $e$ is a self-loop, and 1 otherwise.

RCK infers a multiplicity function $\mu$ that is determined by the underlying cancer genomes, by solving an optimization problem formulated as a mixed integer linear program, which takes into account (i) restrictions that both the previously described evolutionary model and haplotype constraint groups place on which haplotype-labeled versions of measured SVs can be present; (ii) the no-novel-telomeres assumption; (iii) the need to utilize at least a fraction $P \in [0,1]$ of input SVs; and such that the length-weighted segment copy number distance $\|\widetilde{\boldsymbol{C}} - \boldsymbol{C}_{\boldsymbol{\mu}}\|$ between the input allele-specific CNV profile and the one inferred via the edge multiplicity function $\mu$ values on segment edges is minimized, or more simply such the CNV profile of the inferred cancer karyotype is as close as possible to the input one.

We ran RCK v 1.1 on the inferred sample-specific SV callset and both HATCHet and TitanCNA CNV profiles separately with the required fraction $P$ of input SVs to be utilized being set 0.9. We also significantly improved the performance of the original version of RCK by introducing the per-chromosomal pre-processing step. In this step RCK solves the karyotype inference problem on a per chromosome basis, such that the union of solutions would equal to the whole genome problem solution excluding the inter-chromosomal SVs. Per-chromosomal solutions are then used a starting vector for the whole-genome MILP problem solution search. This allowed us to improve performance by reducing the running time from 48 to 6 and from 32 to 8 hours wall clock time for TitanCNA and HATCHet CNV input respectively. RCK was run on a 24 core (with --run-threads 24 flags) machine with 512GB RAM (with a peak usage of ~200GB of RAM). We note that time and high RAM usage is due to tens of thousands of input SVs and haplotype constraint groups; on simpler cancer samples (with ~1000 SVs) RCK can infer cancer karyotypes in several minutes.

Circos plots shown in **Figure 4c**, **Supplementary Figure 3**, **Supplementary Figure 6** and **Supplementary Figure 7** were constructed with Circa v 1.2.0 software (http://omgenomics.com/circa).

**Analysis of COSMIC census genes affected by SVs.**
For the analysis of SVs and COSMIC census gene interactions, we considered genes from the COSMIC Cancer Gene Census v 88.

We considered a COSMIC census gene $g = A: [a, b]$, with a start coordinate $a$ and an end coordinate $b$ on chromosome $A$, being affected by SV $s = \{X: p, Y: q\}$ with breakends $p$ on chromosome $X$ and $q$ on chromosome $Y$ if either $X = A$ and $a \leq p \leq b$ or $Y = A$ and $a \leq q \leq b$, or both. We note that SV's breakends' strand orientations are not important in considering whether a gene is affected by SV, as in either case the breakend disrupts the genomic regions that contains the gene. We say that a COSMIC census gene $g$ is affected by SVs if it is affected by at least one SV, and we say that SV $s$ affects COSMIC census genes if at least one COMSIC census gene $g$ is affected by $s$.

We further analyzed the SVs interactions with the COSMIC census genes with their *flanking sequences* (i.e., for every gene $g = A: [a, b]$, we considered $g_\Delta = A: [a - \Delta, \mathrm{b} + \Delta]$, where $\Delta \in [500, 1000, 5000]$). We did not discover any additional COSMIC census genes affected by SVs nor have we observed any additional SVs affecting COSMIC census genes with any of the considered values of Δ.

**Analysis of COSMIC census genes affected by CNVs.**
For the analysis of CNVs and COSMIC census gene interactions, we considered genes from the COSMIC Cancer Gene Census v 88.

We considered a COSMIC census gene $g = A: [a, b]$, with a start coordinate $a$ and an end coordinate $b$ on chromosome $A$, being affected by an allele-specific deletion (amplification) if there exists a segment $j = A: [c, d]$ that overlaps $g$, or more formally, either $a \leq c \leq b$, or $a \leq d \leq b$, or both, and $j$ has the respective allele-specific CNV: i.e., either $a_j > 1 (< 1)$ or $b_j > 1 (< 1)$. Identification of the COSMIC census genes affected by allele-specific CNVs was performed with the in-house RCK-based utility script. We note that the same COSMIC census gene $g$ (on either the same or different haplotypes) may be simultaneously affected by both allele-specific deletion(s) and amplification(s).

**Complex rearrangements analysis.**
Complex rearrangements reflect an underlying double stranded breakage event of $k \geq 3$ double-stranded breaks. Segments resulting from the breakage are then often amplified or lost and the subsequent ligation of involved segments are then detected as SVs. In general, by simple observation of a group $U$ of SVs we cannot determine whether all of the SVs in $U$ we produced by a single complex rearrangement or by several sequential rearrangements. However, under the considered somatic evolutionary model with the infinite sites assumption we note that any pair $a = \{p^x, j_H^h\}$ and $b = \{(j+1)_H^t, q^y\}$ of *reciprocal SVs* that involve reciprocal extremities $j_H^h$ and $(j+1)_H^t$, which are adjacent in the reference, must be produced by a single rearrangement event which breaks the reference adjacency $\{j_H^h, (j+1)_H^t\}$.

To identify complex rearrangements for a given set $\mathcal{A}$ of SVs we constructed a complex rearrangements graph $G_C = (V, E)$, where a set $V = \left\{\{j^h, (j+1)^t\} \mid \{j_A^h, (j+1)_A^t\} \in \mathcal{A}(R)\right\}$ of vertices is determined by non-unlabeled versions of reference adjacencies, and every edge $e_a = \{u, v\} \in E$ is determined by an SV $a = \{p^x, q^y\}$ such that $p^x \in u$ and $q^y \in v$, or more simply, if $a$ connects extremities involved in $u$ and $v$. Once the $G_C$ is constructed complex rearrangements are determined as connected components with at least 3 vertices in them, as they correspond to groups of reciprocal SVs. We note that not every k-break produces reciprocal SVs, but every group of reciprocal SVs is produced by a k-break. We also note, that haplotype labels are not crucial for determination of complex SVs under the proposed framework. Complex rearrangements inference based on reciprocal SVs grouping was implemented in the RCK v 1.1.

For patient 51 we identified complex rearrangements in both the inferred specific SV set $\mathcal{A}$, as well as in subsets $\mathcal{A}_H \subseteq \mathcal{A}$ and $\mathcal{A}_T \subseteq \mathcal{A}$, which are determined by karyotypes inferences with HATCHet and TitanCNA CNV inputs respectively. We also inferred complex rearrangements signatures on SV subsets $\mathcal{A}_l \subseteq \mathcal{A}$ and $\mathcal{A}_s \subseteq \mathcal{A}$ of the 51T specific SV set as determined by long- and short-read exclusive SVs. For patient 48 and SKBR3 we inferred complex rearrangements on the respective specific SV sets.

### Supplementary Figures

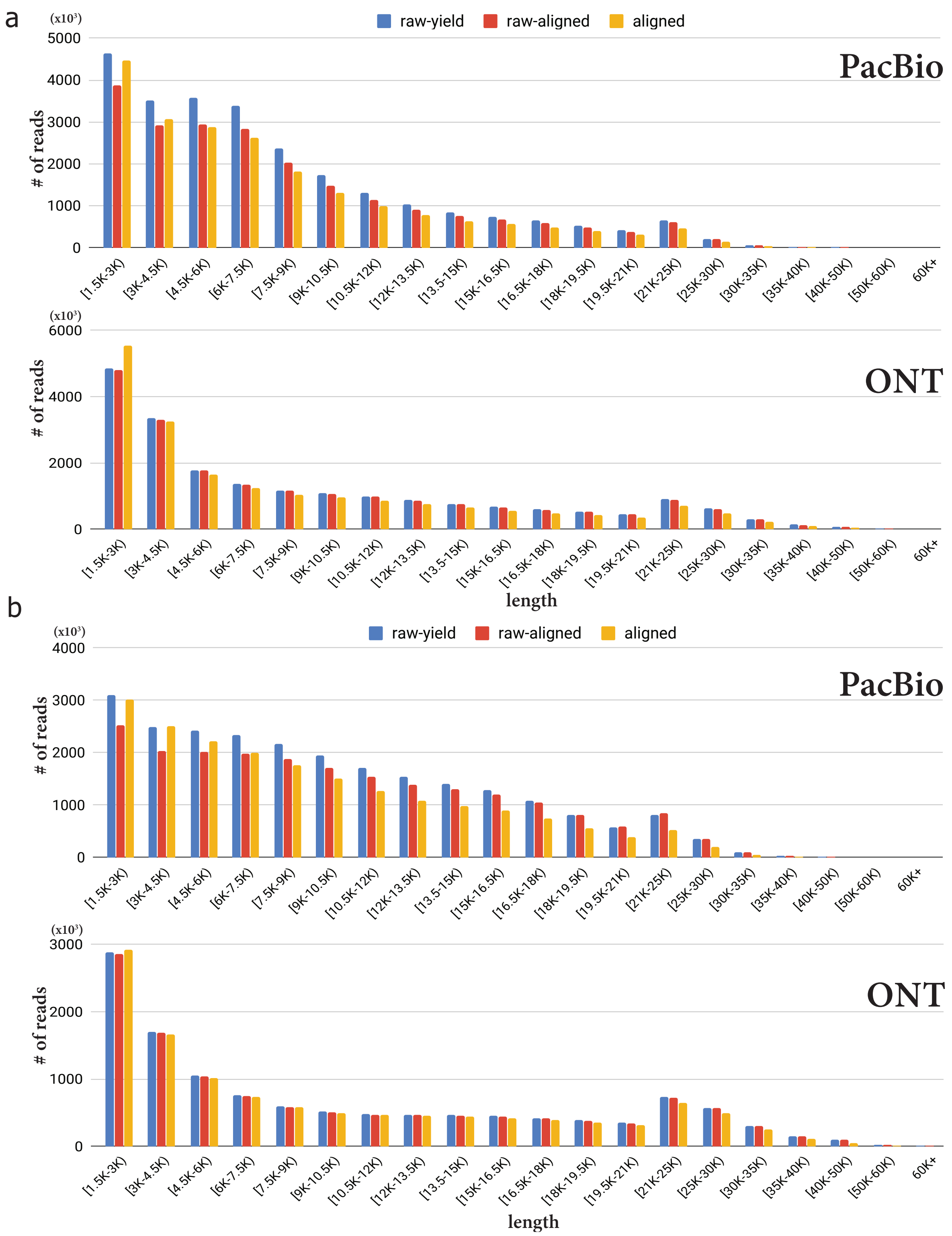


**Supplementary Figure 1. | ONT and PacBio reads lengths distribution.** *raw-yield* corresponds to lengths of raw sequenced reads, *raw-aligned* corresponds to lengths of raw read that had any alignment inferred for them and *aligned* corresponds to lengths of aligned parts of sequenced reads. **a)** Lengths distributions for reads of length *1.5+Kbp* from PacBio and ONT sequencing runs for samples 48T. **b)** Lengths distributions for reads of length *1.5+Kbp* from PacBio and ONT sequencing runs for samples SKBR3.

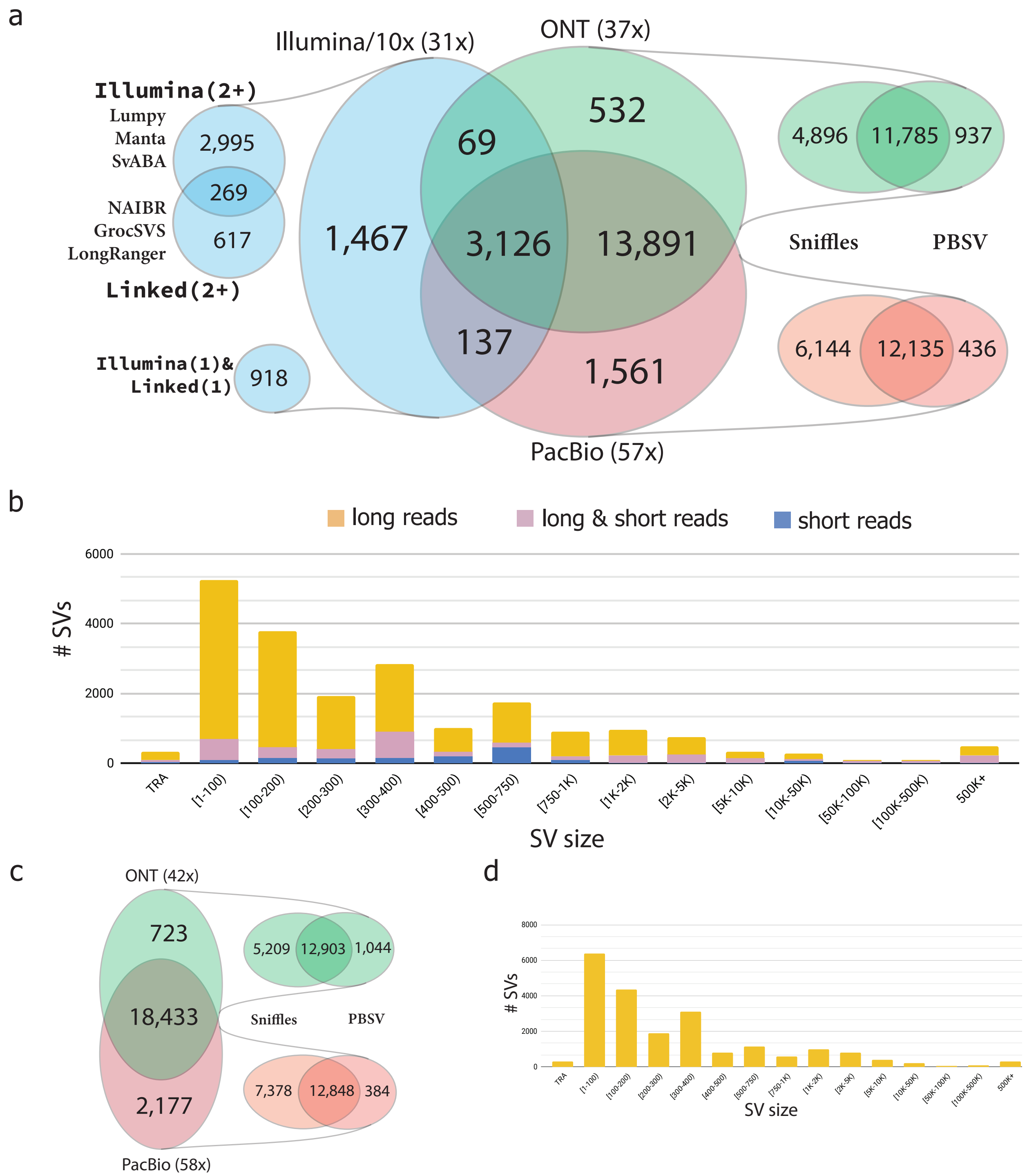


**Supplementary Figure 2. | Structural variantions inference across Illumina/10X, ONT, and PacBio sequencing platforms for samples SKBR3 and 48T. a)** and **c)** SV inference comparison across SVs inferred from *Platform (x)* sequencing exeriments, where *Platform* corresponds to sequencing technology, and *(x)* determines the average alignment read-depth coverage. Methods-specific breakdown is provided for every sequencing technology. **b)** and **d)** Size distribution for SVs in samples SKBR3 and 48Twith SVs being either exclusively inferred from either long-reads (either ONT, or PacBio, or both), or exclusively from Illumina/10x short-reads, or supported by both long and short-reads.

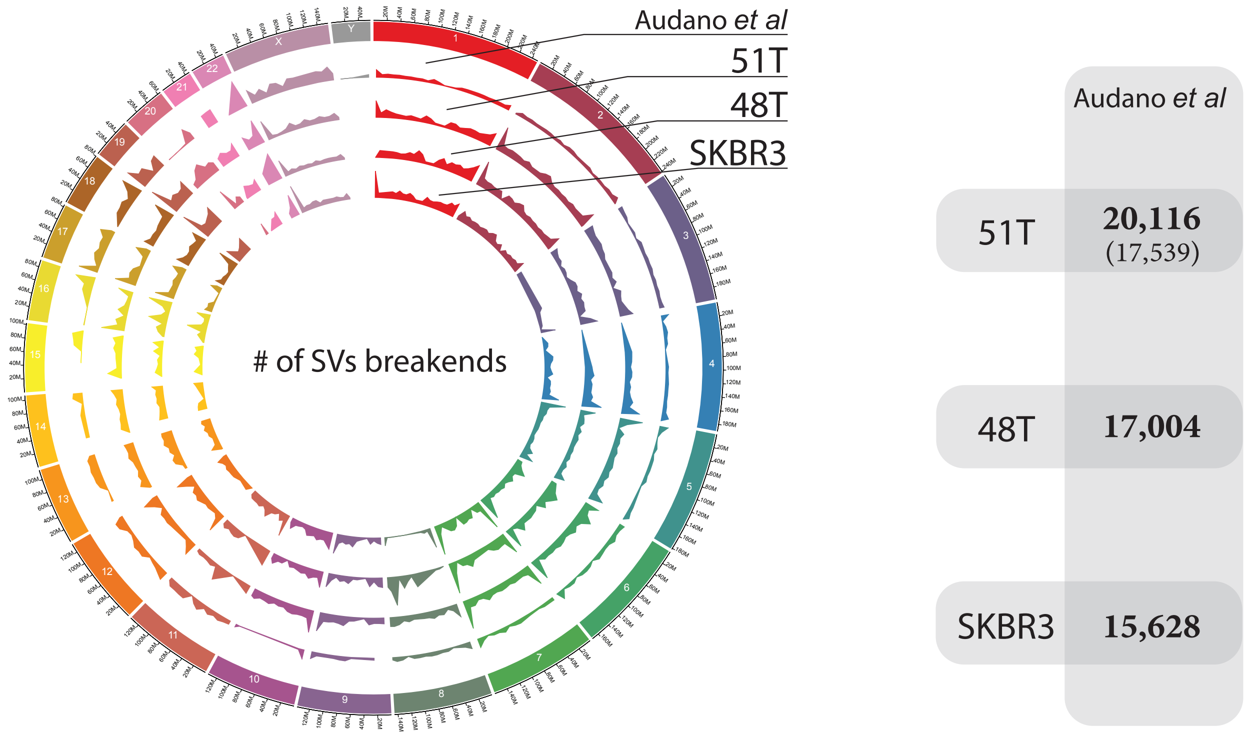


**Supplementary Figure 3. | Structural Variations in samples 51T(N), 48T, SKBR3, and in Audano *et al*[20] dataset.** Circos plot on the left shows the SVs breakends distributions across genome chrmosomes. Every track is dataset-specific shows the total number of SVs' breakends over 5MB segment-length windows. Panel on the right shows intersection of SVs across observed cancer datasets (with matching normal SVs shown in parentheses) and the healthy SV set generated from 15 samples from Audano *et al*.

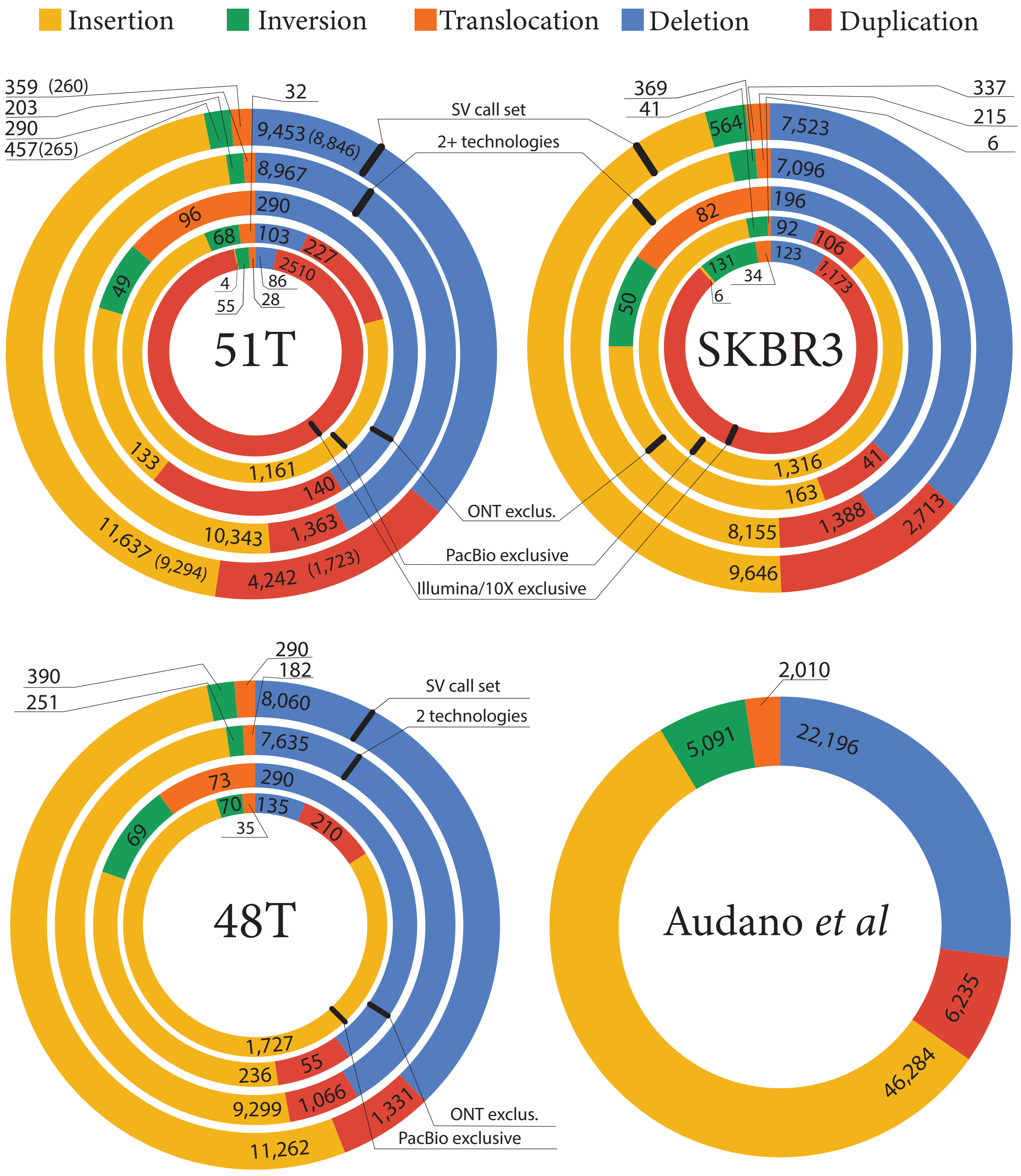


**Supplementary Figure 4.** |**SV type breakdown over supporting sequencing technologies.** Quantification of SVs types determined by involved intra-chromosomal breakends' orientations (*deletion*, *duplication*, *inversion*), inter-chromosomal breakends (*translocation*), and intra-chromosomal SVs with novel sequence (*insertions*) across different combinations of sequencing technologies in samples 51T, SKBR3, 48T, and in the healthy SV set (i.e., the union of SVs across 15 healthy PacBio sequenced samples from the Audano *et al* study dataset).

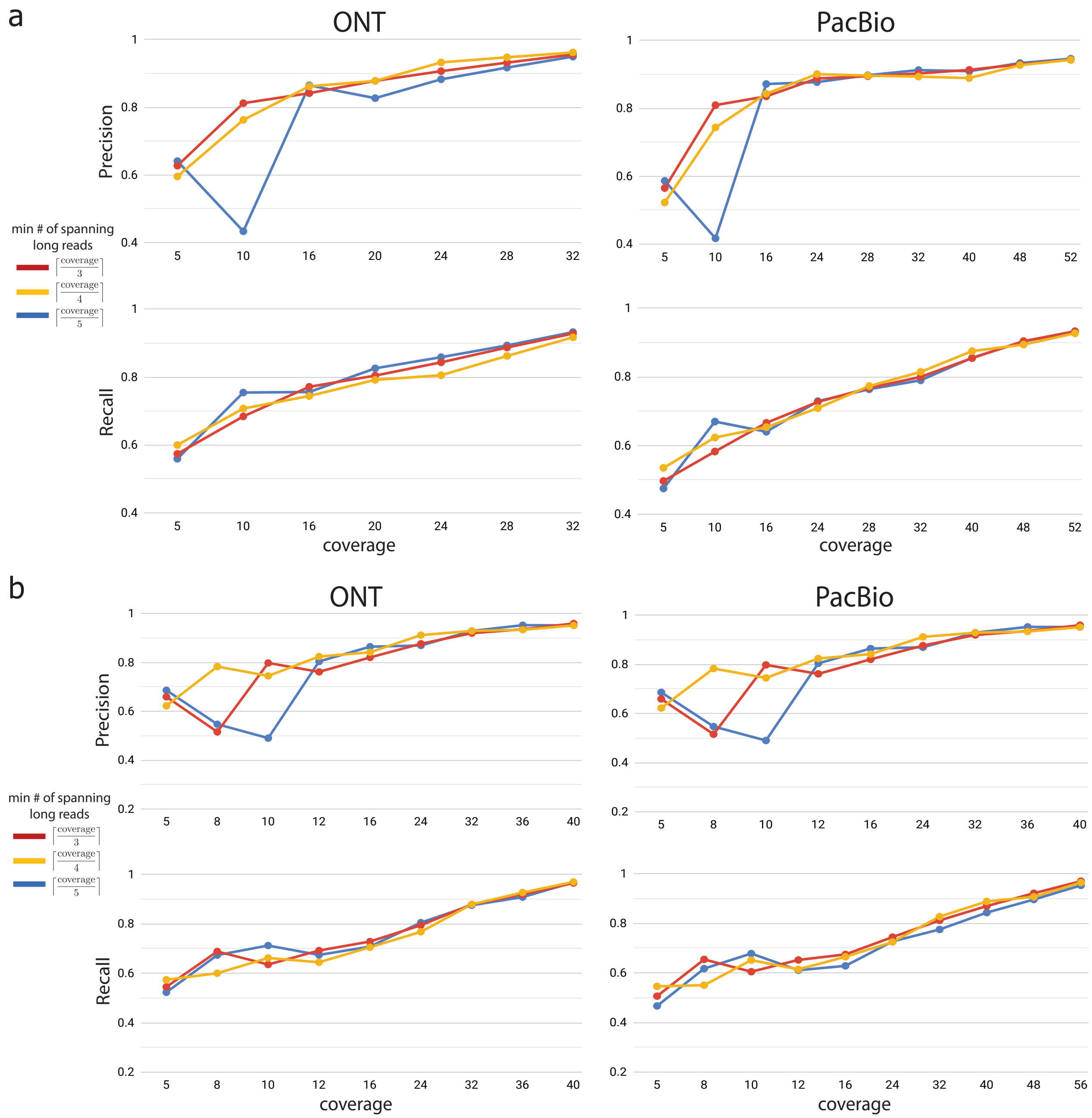


**Supplementary Figure 5. | Concordance between downsampled and full coverage SKBR3 and 48T datasets with distinct minimum fractional *x/y* read support for an SV to be considered. a)** Precision and Recall for SVs inferred on downsampeld ONT and PacBio dataset for sample *SKBR3*, with the SVs inferred on the full coverage dataset used as the ground truth. **b)** Precision and Recall for SVs inferred on downsampeld ONT and PacBio dataset for sample *48T*, with the SVs inferred on the full coverage dataset, at the matching support threshold, used as the ground truth.

a

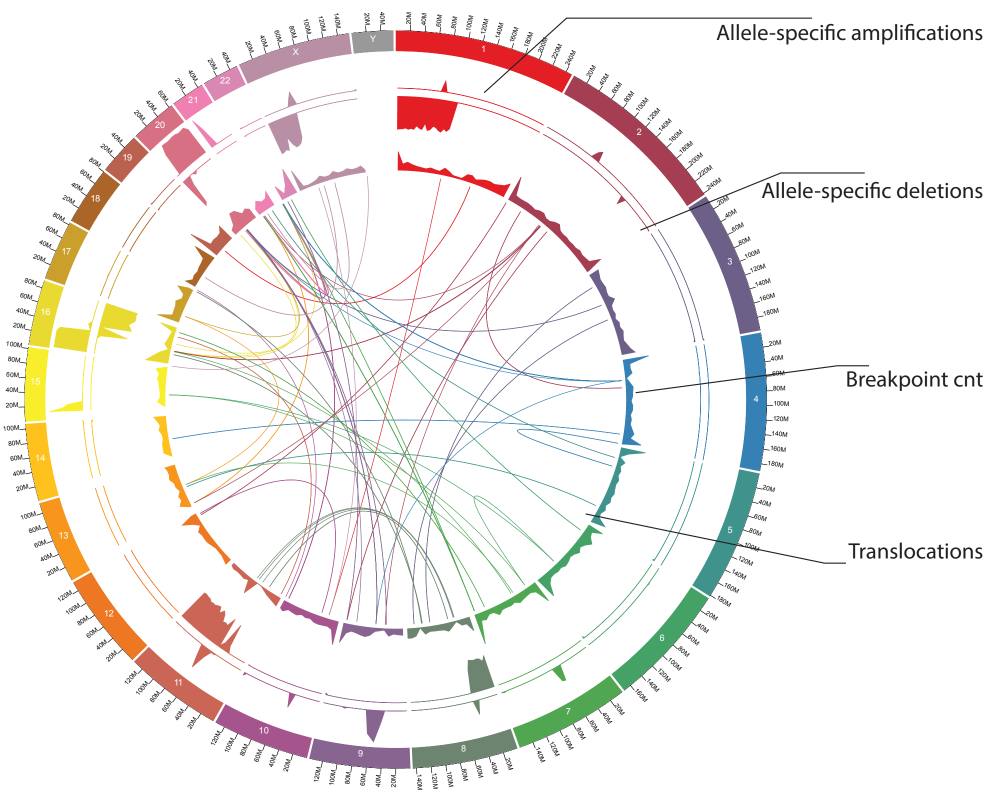


b

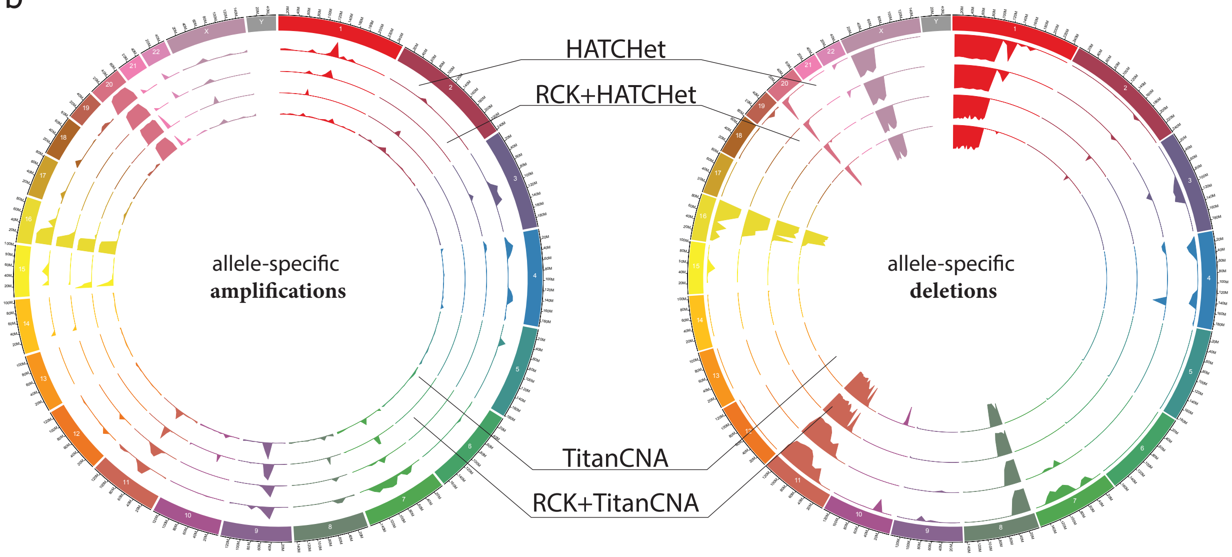


**Supplementary Figure 6.** | **Haplotype-specific karyotype and CNVs comparison for cancer genome in sample 51T**. **a)** Circos plot of the cancer karyotype inferred by RCK for patient 51 with TitanCNA segment copy number (CN) input. Top two tracks corresponding to fractions *x/y* of the total length *x* of either amplified (CN ≥ 1) or deleted (CN = 0) fragments over the $y=5x10^6$ long windows. Breakend track shows the total number (with 590 being the maximum value shown) of breakends inferred by RCK as being present. Translocation track shows inter-chromosomal SVs inferred by RCK as being present in the reconstructed katyorype. **b)** Circos plots of the allele-specific amplifications (left) and deletions (right) across fragments of the $y=5x10^6$ length in raw HATCHet CNV profile (top track), RCK+HATCHet karyotype (second track), RCK+TitanCNA karyotype (third track), and raw TitanCNA CNV profile (fourth track).

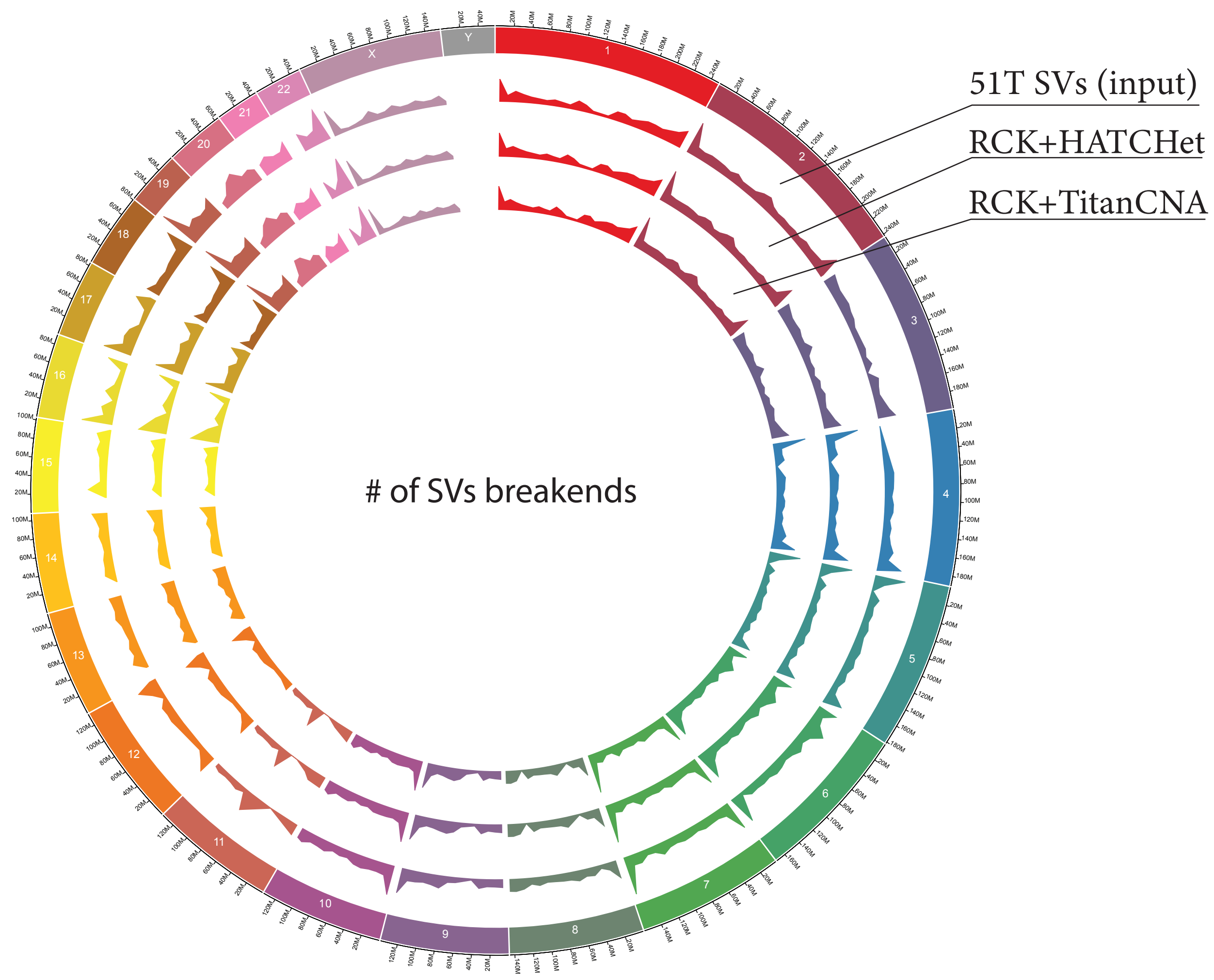


**Supplementary Figure 7. | Breakpoints distribution in both raw and karyotype-utilized SV callsets for sample 51T.** Circos plot of the SVs breakends distributions across genome chrmosomes. Every track is dataset-specific shows the total number of SVs' breakends over 5MBp segment-length windows in the 51T specific SV callset (top track), and SVs utilized in the karyotype reconstrctured by RCK with HATCHet (second track) and TitanCNA (third track) allele-specific CNVs input.

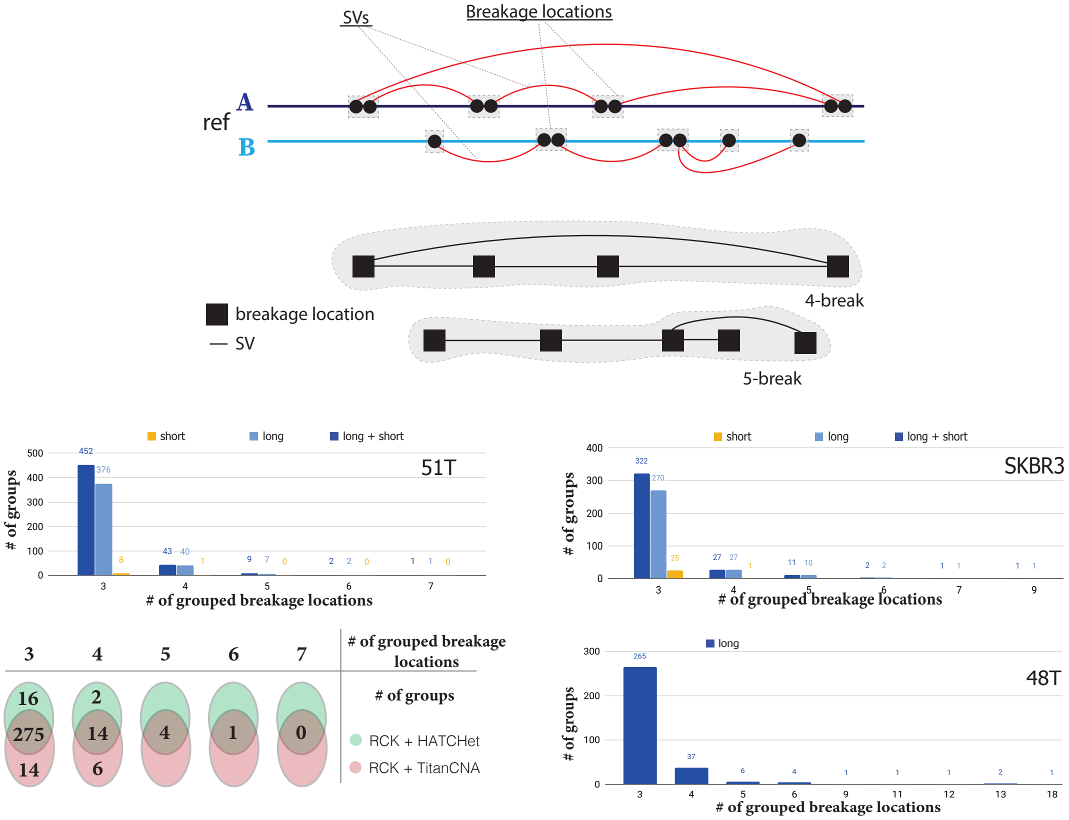


**Supplementary Figure 8. | Complex Rearrangments in 51T, SKBR3 and 48T cancer samples.** (Top) Signature of two complex k-break rearrangements (i.e., 4- and 5-breaks) as evident by breakage locations linked via reciprocal SVs. Statistics over k-break rearrangments identified in the input SV callset for samples *51T, SKBR3, and 48T* (with a breakdown over long, short, and long+short-read support) and in RCK+HATCHet and RCK+TitanCNA karyotypes for sample 51T.

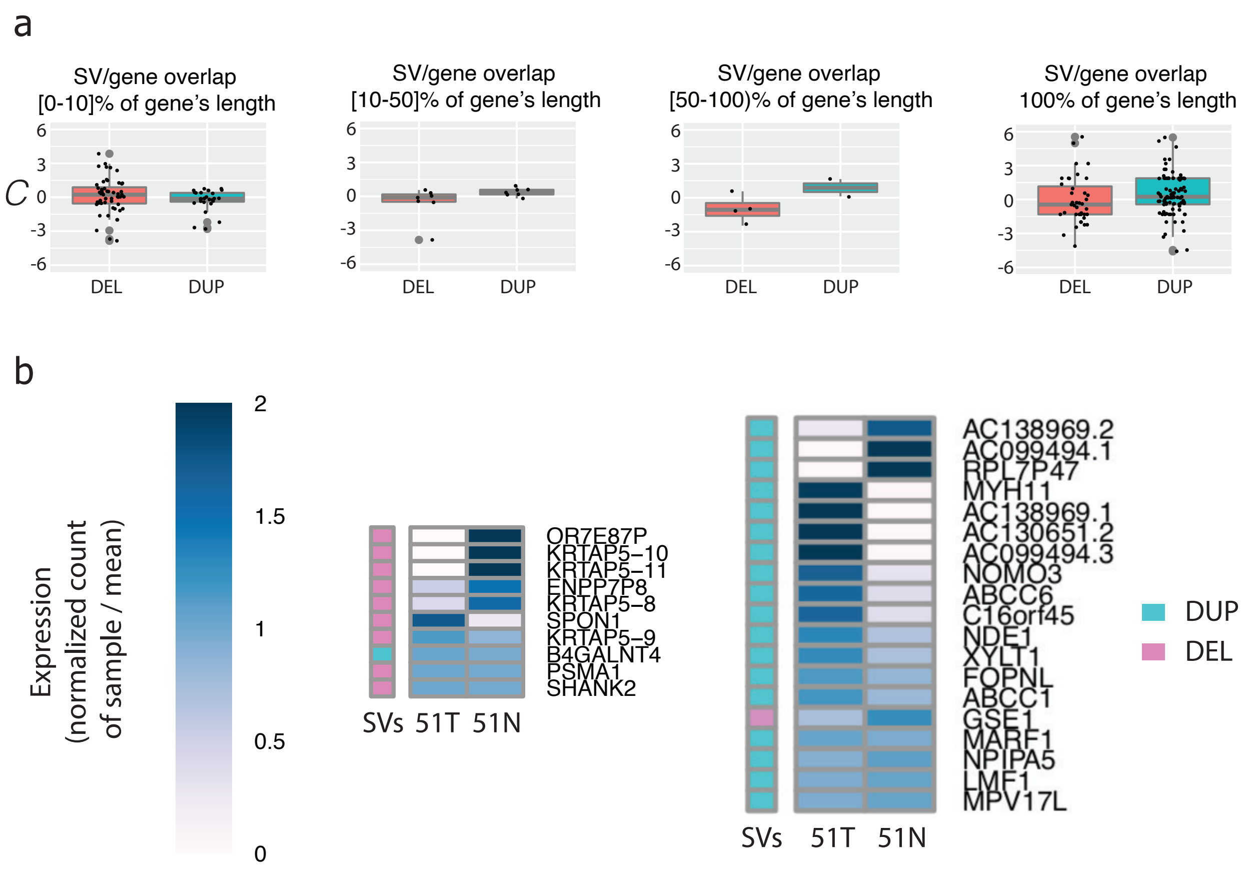


**Supplementary Figure 9. | RNA-seq expression analysis in patient 51 for genes overlapped by SVs present only in sample 51T and not in 51N and not in the union of SVs from 15 healthy genomes. a)** Changes in expression for genes overlapped by deletion and duplication SVs quantified by the percentage of genes' lengths spanned by SVs. Every gene$g$ is represented as a dot, with $C(g) = \log_2(T(g) + 0.5) - \log_2(N(g) + 0.5)$, where $T(g)$, and $N(g)$ are expression count for $g$ in 51T and 51N samples respectively. b) Groups of genes affected by Deletions and Duplications. **b)** examples of long DUP/DEL SVs spanning multiple genes and changes in respective genes' expression levels in 51T vs 51N samples respectively.

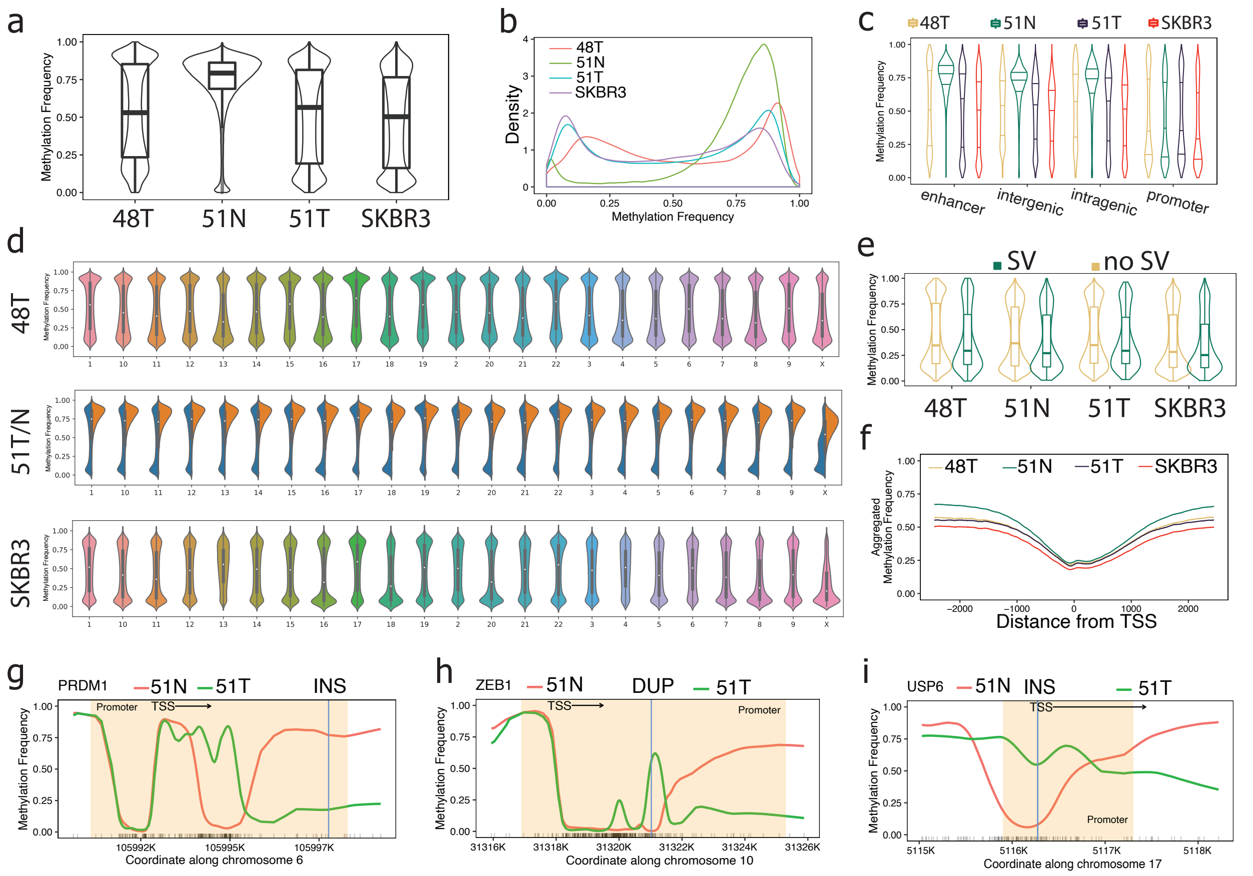


**Supplementary Figure 10. |Methylation analysis on cancer samples 48T, 51N, 51T, and SKBR3. a)** Genome-wide distribution of methylation frequencies. **b)** Genome-wide methylation frequency density functions. **c)** Methylation frequency distributions over 1Kbp windows that overlap enhancer, intergenic, intragenic, and promoter regions respectively. **d)** Per-chromosomal methylation frequency distributions. **e)** Methylation frequency distributions in the 1Kbp windows which overlap promoter regions and either contain or do not contain SVs breakends respectively. **f)** Averaged aggregated methylation frequencies around Transcription Start Sites (TSS). **g)** Methylation frequencies in PRDM1 gene promoter region with transition between hypermethylation in 51N and hypomethylation in 51T around identified insertion. **h)** Methylation frequencies in ZEB1 promoter region with hypermethylated region in 51T coinciding with identified duplication SV. **i)** Methylation frequencies in USP6 gene promoter region with identified insertion coinciding with blocking of the TSS from demethylation in 51T.
